# During natural vision, semantic novelty modulates fixation-related processing in primate cortex

**DOI:** 10.64898/2026.03.18.712708

**Authors:** Vinay S Raghavan, Jens Madsen, Maximilian Nentwich, Marcin Leszczynski, Arnaud Falchier, Stephan Bickel, Brian E. Russ, Lucas C. Parra

## Abstract

We sample visual scenes with short gaze fixations separated by saccades. While low-level integration is known, semantic integration of foveal vision across fixations remains unclear. We hypothesized that the brain responds to changes in semantic information from one fixation to the next, and therefore postulated a neural signal associated with semantic novelty for each saccade. Novelty was measured from different layers of a deep network on foveal vision. Only semantic, and not low- or mid-level, novelty modulated frontal and occipital fixation-related potentials in male and female human scalp EEG during natural viewing of full-length movies (3.4×10^6^ saccades). Intracranial recordings in male and female humans (9.0×10^4^ saccades) and female non-human primates (Macaca mulatta; 3.3×10^4^ saccades) revealed broadband high-frequency activity modulations in scene-specific ventral visual and frontal brain areas only for semantic novelty. This modulation was stronger for movies than static images, and frontal modulation occurred early alongside occipital modulation, suggesting top-down effects. This modulation of fixation-related activity with novelty suggests that foveal representations are integrated across fixations to construct scene representations during natural viewing in primates.

**Significance statement:** Primate vision relies on eye movements to sample the world, yet how the brain integrates high-level meaning across these “snapshots” remains a mystery. By combining deep-learning models with massive electrophysiological datasets from humans and macaques, we show that the brain responds to changes in semantic content between fixations, but not low- or mid-level changes. We identified a “semantic novelty” signal that modulates neural activity across the primate cortex. These findings suggest visual processing is not a series of independent glimpses, but a continuous process integrating foveal information at the semantic level. This work bridges biological vision and artificial intelligence, providing a new framework for understanding how the brain maintains a coherent understanding of a dynamic world.

## Introduction

Humans and other animals shift gaze via saccades followed by stable fixations to actively sample the visual environment with high-resolution foveal vision (Robinson, 2022). These eye movements evoke strong neural responses previously measured with scalp recordings (Kazai and Yagi, 1999; Degno and Liversedge, 2020) and intracranial recordings in humans (Bartlett et al., 2011; Barczak et al., 2019; Leszczynski et al., 2021, 2023) and non-human primates (Rajkai et al., 2008).

Prior evidence points to the integration of visual information across saccades. Peripheral input first aids target selection (Deubel and Schneider, 1996) and is then integrated with foveal input to maintain a continuous experience (Irwin and Gordon, 1998; Kämmer et al., 2026; Kroell and Rolfs, 2026). Integration between foveal and peripheral vision occurs for attended, low-level features and object sizes (Wolf and Schütz, 2015; Valsecchi and Gegenfurtner, 2016). Whether foveal information is integrated across fixations remains unknown. Low-level integration may simply be expected from a change in visual input associated with eye movements, while high-level foveal-to-foveal integration would help construct a coherent scene representation from separate fixations. Semantic information plays a role in directing eye movements in static and dynamic scenes (Eisenberg and Zacks, 2016; Henderson and Hayes, 2017). Nevertheless, it remains unclear whether the brain integrates semantic visual information across consecutive fixations.

During a single fixation, the ventral visual stream extracts semantic information through a sequence of visual processing areas (Logothetis and Sheinberg, 1996), producing representations similar to deep vision models (Yamins et al., 2014; Yamins and DiCarlo, 2016; Lindsay, 2021). However, these computer vision models differ from biological vision in two key ways: they are limited to a single fixation, rather than multiple fixations, and they are often trained using supervised methods, i.e., with hand-labeled images.

Meanwhile, a revolution in AI has emerged avoiding training labels with self-supervised learning techniques compatible with natural vision. One particularly effective method learns robust representations by predicting the embedding of one crop of an image from another (Chen and He, 2021), comparable to predicting the semantic representation of one fixation from the last.

The embeddings learned by these deep networks capture semantic content by directly comparing representations between crops of the same image (Wen and Li, 2021). These representations are considered semantic since they are linearly related to distinct object categories, which are distinct semantic concepts (Reilly et al., 2025).

We hypothesized a similar mechanism is at play in primate vision, whereby the semantic content of foveal vision is compared across fixations. If so, we expect semantic novelty alone to modulate neural activity associated with fixations while viewing static and dynamic scenes, with no modulations due to low- or mid-level novelty. We defined novelty features as the distance between embeddings of successive foveal fixation patches extracted from different layers of a deep network (Chen and He, 2021).

We extracted novelty from 4 hierarchical stages of the deep network, representing multiple scales of hierarchical novelty. We find that more common low- and mid-level visual features, as well as salience features, are uncorrelated with these novelty features (Fig S1). Additionally, we are interested in how the brain responds to the integration of foveal content, rather than predicting saccade targets or modeling saccade control (Schiller and Tehovnik, 2005; Kümmerer and Bethge, 2023).Thus, we only modeled eye movement onsets, saccade amplitude, and hierarchical novelty features. We used cross-validated linear encoding models that account for correlated and delayed responses that may overlap between adjacent eye movements (Crosse et al., 2016; Holdgraf et al., 2017; Ehinger and Dimigen, 2019).

Our findings in human scalp potentials and intracranial recordings in human and non-human primates demonstrate a key role for semantic novelty alone in modulating fixation-related neural responses. By rigorously controlling for low- and mid-level novelty features, we isolate a semantic novelty response that could build scene representations across eye movements and tune the ventral visual system. This work provides evidence that semantic information is integrated between fixations.

## Materials and Methods

### Participant details

The human scalp dataset comprised 79 participants (41 female), aged 18-37 years (M = 23.43, SD = 4.67), whose data met inclusion criteria and were included in the analyses. All procedures were approved by the Institutional Review Boards of the City University of New York, and all participants provided informed consent.

The human intracranial dataset comprised 31 patients (age 18-59, mean 38, 16 female) with a total of 8522 electrodes implanted. All patients are included in the image dataset, and a subset of 23 patients (6328 electrodes) is included in the movie dataset (age 19-58, mean 38, 11 female). Patients were implanted with depth, grid, and/or strip electrodes for clinical treatment of drug-resistant epilepsy at Northwell Health (NY, USA). Four patients were implanted twice at different times, and sessions were recorded again from these patients to capture the new electrode coverage. Three re-implants are included in the movie dataset. The study was approved by the institutional review board at the Feinstein Institute for Medical Research, all clinical investigation was conducted according to the principles expressed in the Declaration of Helsinki, and all patients gave written informed consent before electrode implantation.

The NHP dataset included 2 rhesus macaques (Macaca mulatta; 2 female, age 4-9). All procedures were approved by the Institutional Animal Care and Use Committee of the Nathan Kline Institute and were carried out in accordance with NIH standards for work involving non-human primates. The two animals, NHP T and NHP W had a total of 27 and 23 electrodes implanted in cortical or subcortical regions, respectively.

### New versus existing data

In total, judging by duration, 95.2% of the data in this study have not been previously analyzed. The fraction of the human iEEG data that have been previously analyzed (79.7%) did not study fixation-related responses for visual features as we have done here. Specifically, the four distinct datasets are as follows: New human scalp data with movies (238 viewings, 22,000 min total). Existing human intracranial data with movies (23 patients, 1063.7min total, with 23 previously analyzed in (Nentwich et al., 2023); 22 in (Nentwich et al., 2025); 21 in (Mishra et al., 2025); 8 in (Leszczynski et al., 2023)), largely new human intracranial recordings with images (9 of 31 patients, 93.6min of 387.6min previously analyzed in (Leszczynski et al., 2023)), and new non-human primate intracranial with movies and images (movies 133min+73min; images: 56min + 13min). Separate data release papers are forthcoming for these datasets.

### Experimental materials and procedure

Each participant in the human scalp movie dataset watched between 1 and 10 full-length movies (1.5–2 h) selected from the following set: *The Big Sick, The Peanut Butter Falcon, Whiplash, Room, Me and Earl and the Dying Girl, The Tomorrow Man, Dom Hemingway, Life After Beth, Woodshock,* and *The Comedian*. After applying exclusion criteria, 79 participants contributed a total of 244 participant–movie viewings across these 10 movies, with participants watching an average of 3.09 movies (median = 2; range = 1–10). Participants were positioned in a comfortable reclinable chair with a neck rest and wore a headrest to constrain head movement while viewing a screen subtending approximately 32.2 degrees horizontally and 19.5 degrees vertically, with each movie shown at 720×404 filling the screen both horizontally and vertically. Participants were instructed to watch the movie as they would normally watch a movie. After viewing, participants answered questions about the movie (not used in this study).

Each participant in the human intracranial movie dataset watched 29.3 to 43.7 minutes of videos (mean ± sd: 40.9 ± 4.7 minutes), for 1063.7 minutes total. The videos included a 10 minute clip of the animated movie Despicable Me (English-language), a non-overlapping 10 minute clip of Despicable Me (Hungarian-language), a 4.3 minute clip of the animated short film The Present (English-language), and three 5 minute clips of documentaries of macaques (no sound), all used previously (Russ and Leopold, 2015; Russ et al., 2021; Nentwich et al., 2023; Telesford et al., 2023). Each participant in the human intracranial image dataset viewed 6.1 to 28.1 minutes of images (mean ± sd: 12.5 ± 3.7 minutes) for a total of 387.6 minutes. 80 unique images were presented, each viewed no more than once. The image content ranged from animals, people, and food to animated movies, vehicles, and toys. Participants in both movie and image human iEEG datasets viewed a display subtending approximately 40.0 degrees horizontally and 25.3 degrees vertically. Movies were presented at a resolution of 1024×768, filled the screen vertically, and filled three-quarters of the screen horizontally, due to differing aspect ratios.

Images were presented at a resolution of 1536×864 with a black border taking up 10% of the screen.

Each NHP watched the same video clips and viewed the same images as the human intracranial participants. Each video was watched multiple times by each NHP. NHP T watched each of the three macaque documentaries 4, 5, and 5 times, for a total of 133 minutes. NHP W watched each of the three macaque documentaries 3, 5, and 5 times, and each of the other videos twice each (Despicable Me - English, Despicable Me - Hungarian, The Present), for a total of 73 minutes. NHP T and W both viewed each image several times, for totals of 56 and 13 minutes, respectively. Both NHPs viewed a display subtending approximately 25.2 degrees horizontally and 19.0 degrees vertically. Both movies and images were presented at a resolution of 1024×768 filling the screen both horizontally and vertically.

### Data acquisition

Human scalp EEG was recorded at 2048 Hz using a BioSemi ActiveTwo system with 64 scalp electrodes placed according to the 10-10 international system. Four electrooculogram (EOG) electrodes were placed above, below, and lateral to the eyes to capture ocular artifacts. Eye position (single eye) was recorded using the SR Research EyeLink 1000 system with a 35-mm lens at a sampling rate of 500 Hz allowing free head movements. A standard 9-point calibration was followed by manual verification. Head movement was constrained by subjects wearing a headrest as they sat in a reclined comfortable chair. Saccades and blinks were detected using the device’s built-in algorithm.

Intracranial EEG from human participants was recorded from stereoelectroencephalography depth electrodes, subdural grids, and/or subdural strips (Ad-Tech Medical Instrument Corp., Oak Creek, WI, USA; Integra LifeSciences, Princeton, NJ, USA; PMT Corp., Chanhassen, MN, USA). Subdural grid/strip contacts were 3-mm platinum disks with 10-mm intercontact spacing. Depth electrode contacts were 2-mm cylinders with 0.8-mm diameter and 4.4- or 2.2-mm intercontact spacing. Intracranial electrode signals were referenced to a subdermal electrode or subdural strip. Data were sampled at 3 kHz (16-bit precision, range ± 8 mv, DC) on a Tucker-Davis Technologies data processor (TDT, Alachua, FL, USA), or at 1KHz an XLTEK Quantum Amplifier (Natus Medical) for 2 of the patients in the image dataset. Gaze position from both eyes was recorded simultaneously with a Tobii TX300 eye tracker (Tobii Technology, Stockholm, Sweden) at 300 Hz allowing free head movements. The eye tracker was recalibrated after each video to prevent drift. Parallel port triggers were sent from the stimulus PC to the eye tracker and the data processor to align the data streams. Custom scripts for movie and image presentation with Psychtoolbox (version PTB_Beta2014-10-19_V3.0.12; Gstreamer version 1.10.2) and for collecting eye tracking data with the Tobii SDK were implemented in MATLAB (2012b, Windows 7). For additional accuracy in the alignment of streams, we recorded timestamps at the onset of each video frame with the clock of the eye tracker.

All procedures were approved by the IACUC of the Nathan Kline Institute for Psychiatric Research. Two female rhesus macaque (*macaca mulatta*) monkeys (at time of recordings: NHP-T: 4.5 yrs ∼4.5-5.25 kg; NHP-W 9 yrs, 4.5-6.0 kg) participated in the current experimental protocol. All experimental tasks were run using SR Research Experiment Builder (SR Research Ltd., Mississauga, Canada).

Prior to neural recordings, animals were implanted with MRI-compatible headposts and trained to fixate visual targets for reward, in order to calibrate their eye positions for future studies. Both animals were then implanted with Spencer sEEG electrodes (Ad-tech, USA) each with 12 contacts (5 mm spacing) covering one hemisphere of the brain. We obtained structural MRIs (Siemens Trio 3T: 4-6 MPRAGE T1w images) prior to implantation, which were used to target the positioning of the sEEG probes. Electrodes were implanted through a craniotomy over the occipital lobe and extended toward the anterior portion of the brain using a stylet till they reached their target location. The first animal, NHP-T, was implanted with three 12 channel sEEG probes extending from the occipital lobe (V1) towards the midbrain (putamen), the anterior ventral temporal cortex (area TE), and the frontal lobe (area 13a&b). The second animal, NHP-W, was implanted with four 12 channel sEEG probes extending from the occipital lobe (V1) towards the superior temporal cortex (rostromedial cortex), the anterior ventral temporal cortex (area TEm), the midbrain (LGN), and the frontal lobe (striatum). Following implantation, the final electrode positions were confirmed with a second MRI imaging session.

Following a 6-8 week recovery period, we began recording neural data while the animals participated in a free-viewing paradigm. Each session, the animals performed a 3 point calibration. Eye position was sampled using EyeLink 1000 system (SR Research Ltd., Mississauga, Canada) at 1000Hz. After calibration the animals participated in blocks of movie viewing, where they were allowed to freely view 4-10min movies. We recorded local field potentials during movie watching using a NeurOne EEG system (Bittium, Oulu, Finland) at 40kHz.

### Intracranial electrode localization

In the human intracranial dataset, the electrode arrays each contained multiple contacts, which were identified using the iELVis MATLAB toolbox (Groppe et al., 2017). All participants received a preoperative T1-weighted 1mm isometric scan on a 3T scanner. Tissue segmentation and reconstruction of the pial surface were performed with the FreeSurfer package. Postoperative CT scans were acquired and coregistered to the FreeSurfer reconstruction. Contacts were semi-manually localized using the BioImage Suite (version 3.01)(Papademetris et al., 2006). All contacts were then coregistered to the FSAverage brain for visualization and assignment to anatomical atlases (see *Human intracranial EEG brain plots*).

In the NHP intracranial dataset, depth electrodes were visualized with a set of post-operative T1-weighted .6mm isotropic scans. The images were registered to each other using AFNI’s 3dAlineate command (Cox, 1996), averaged, and aligned to NMT Atlas space (Seidlitz et al., 2018; Jung et al., 2021). Contact locations were then determined based on their location relative to the CHARM and SARM hierarchical atlases (Seidlitz et al., 2018; Hartig et al., 2021; Jung et al., 2021).

### Neural data pre-processing

The human scalp EEG data were band-pass filtered (0.5-64 Hz), downsampled to 100 Hz, and manually inspected for noisy channels. Interpolation of bad channels was conducted using neighboring electrodes in a 3D-projected coordinate space. Robust Principal Component Analysis (Robust PCA) was applied for artifact removal in both EEG and EOG channels. Eye-movement artifacts were further removed via linear regression of the EOG channels from the EEG data. Residual outliers exceeding ±4 interquartile ranges were replaced by interpolation using neighboring electrodes within ±40 ms.

Both human and NHP intracranial datasets had the same preprocessing steps, all implemented using naplib-python (Mischler et al., 2023). The raw data were resampled to 600 Hz, and a 4th-order Butterworth high-pass filter with a cutoff of 0.5 Hz was used to remove DC drift. Data were re-referenced using a local average scheme whereby each electrode was referenced relative to the average of its nearest neighbors, which depend on the physical configuration of each electrode array. Line noise at 60 and 120 Hz was removed with order 501 FIR filters with 1Hz bandwidth applied forwards and backwards for zero phase shift. The data were then filtered into the high gamma band/broadband high-frequency (70-150 Hz), which reflects local population activity that is related to, but dissociable from, neuronal spiking (Leszczyński et al., 2020). To obtain the broadband high-frequency amplitude (BHA), we used the naplib-python toolbox to filter the data into 8 frequency bins logarithmically spaced between 70 and 150 Hz using Chebyshev Type 2 filters. The envelope of each band was obtained by taking the magnitude of the analytic signal obtained via the Hilbert transform. All eight frequency band envelopes were summed, and the resulting signals were resampled to 100 Hz and z-scored.

### Eye movement detection

In the human scalp and NHP intracranial datasets, eye movements were recorded with SR Research EyeLink hardware, and the default parameters were used for eye movement detection. Specifically, saccades were detected as periods in which angular velocity and acceleration were greater than 30°/s and 8000°/s^2, respectively.

In the human intracranial datasets, eye movements were recorded with the Tobii TX300, so additional pre-processing was needed to detect saccades. A 20th-order median filter was used to smooth gaze position data. This reduces high frequency noise that can lead to false positive saccades. Angular velocity was computed for easier interpretation of saccade amplitude.

Saccades were labeled as samples of eye velocity greater than 2 standard deviations from the average. Short adjustments of gaze position after the saccade are merged into the saccade using a morphological closing operation with a kernel size of 5 samples so that the overshoot is not detected as a separate saccade. The fixation onset corresponds to the first sample after which eye velocity drops under the 70th percentile, computed from velocity values within 33 ms before and 120 ms after saccade onset. The eye tracker provides labels for data quality when the gaze was not detected, e.g., during eye blinks. Saccades within 83 ms of samples with low data quality are removed, because this time period included gaze adjustments after blinks We also remove saccades following fixations less than 110 ms long, which are on the lower range of typically observed intersaccade intervals (Otero-Millan et al., 2008).

Saccade amplitude was calculated for all datasets by first determining the change in horizontal (x) and vertical (y) gaze angles between the saccade’s onset and offset. The amplitude was then defined as the 2-norm (Euclidean distance) of this change vector.

### Fixation patch and feature extraction

For each fixation, we extracted an image patch subtending 5 degrees of the visual field, corresponding to the size of the fovea anatomically (Kroell and Rolfs, 2026). This also matches previous related work (Nentwich et al., 2023; Madison et al., 2025). This offered a balance between providing semantic information and limiting the size to high-resolution vision. A patch size of 5 degrees corresponds to different patch resolutions depending on the physical position of the participant and monitor. In the human scalp, human intracranial, and NHP intracranial datasets, patch resolutions were defined as 100×100, 200×200, and 200×200, respectively. These were computed based on the screen resolution, screen size, and distance between the participant and the screen.

We extracted 37 distinct features related to each fixation patch to test if any of them correlated with any hierarchical novelty features (Fig. S1). In addition to the feature value at each patch, we also measure the difference in feature value between fixations, analogous to how novelty features are measured. From the CIE L*a*b* representation of the image patches, we computed average luminance, luminance skew, luminance kurtosis, average a* channel color, and average b* channel color. From the HSV representation, we computed average saturation and hue diversity. From the grayscale image, we computed the Michelson contrast, RMS contrast, sharpness, vertical energy, horizontal energy, orientation bias, spectrum slope, global entropy, average local entropy with a 7×7 kernel, and energy in the histogram of oriented gradients. We used the grayscale image to extract additional texture and shape features, including local binary pattern energy, Scharr edge density, Sobel edge density, Hough line count, and Hough circle count. We also extracted gray-level co-occurance matrix (GLCM) features, including contrast, homogeneity, correlation, and angular second moment. We defined blobs by thresholding each patch at the mean grayscale value and performed blob analysis, computing the number of blobs, average eccentricity, and average solidity. Finally, we computed salience features, including the intensity (6 maps), color (6 RG, 6 BY maps), and orientation (4 orientations, 6 scales) salience features described previously, as well as DeepGaze III saliency of the fixated location in the 100 to 400ms preceding saccade onset (Itti and Koch, 2000; Kümmerer et al., 2022). We also extracted optical flow features (Nentwich et al., 2023; Yao et al., 2024).

Hierarchical novelty features were computed using a convolutional neural network trained using the Simple Siamese representation learning algorithm (SimSiam)(Chen and He, 2021). In all datasets, fixation patches were upsampled to 224×224 for input to the network using 3rd order spline interpolation. Representational embeddings of the pre- and post-saccadic patches were extracted from 4 main layers of a ResNet50 trained with SimSiam, specifically, following the conv2_x, conv3_x, conv4_x, and conv5_x layers, as indicated in Fig 1B. Novelty is defined as the cosine distance between these embeddings. A large cosine distance between embeddings corresponds to unrelated image patches and thus, a high novelty. Novelty features were not computed for saccades crossing film cuts. Beyond using a SimSiam ResNet50 and extracting novelty from several layers, this process was otherwise similar to previous work (Nentwich et al., 2023).

**Figure 1.**
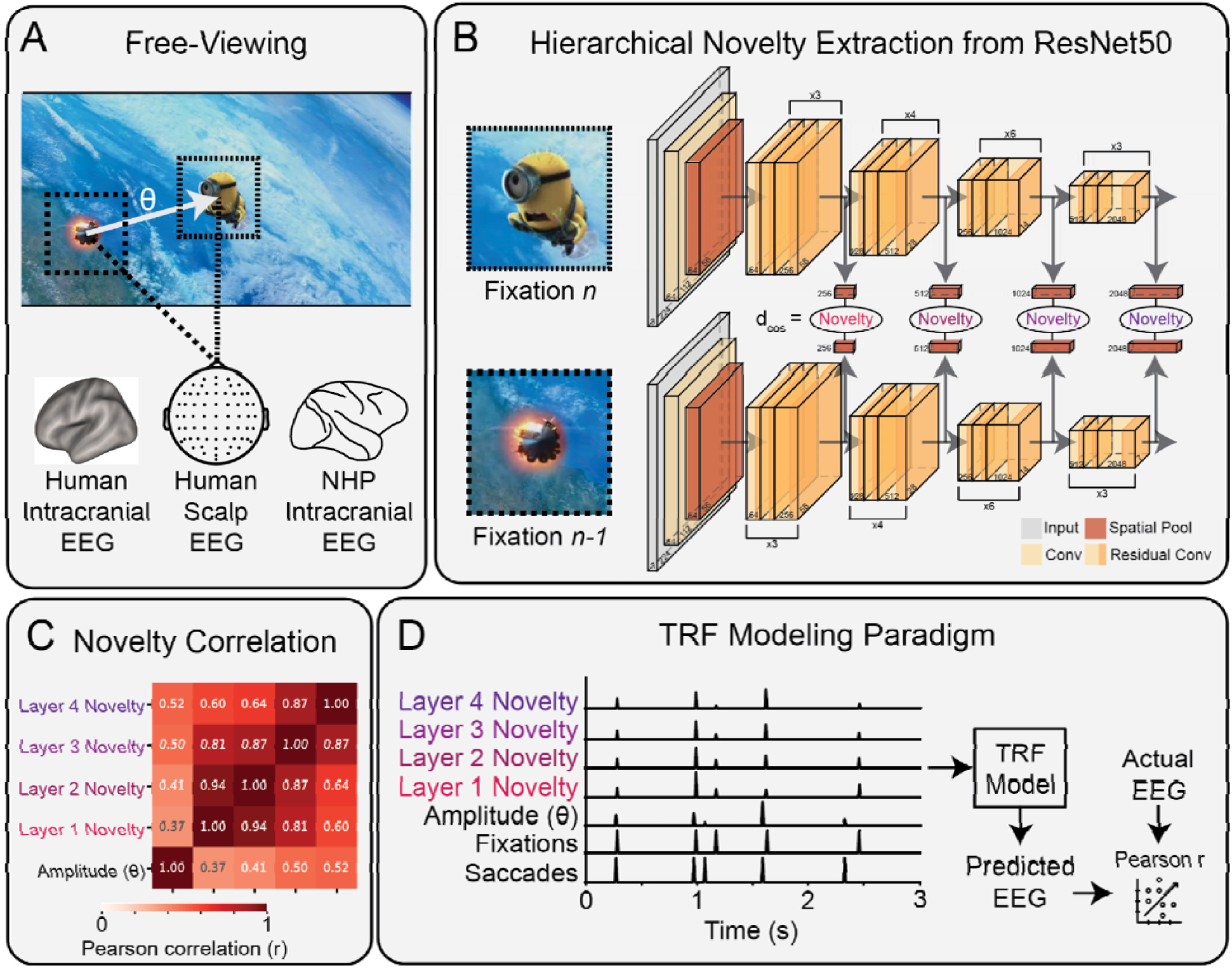
Examples of novelty computation and modeling paradigm. A) Illustration of a saccade, the amplitude (θ), and the fixation patches (5°) before and after the saccade, as measured alongside human scalp, human intracranial and NHP intracranial EEG. B) Novelty is defined as the cosine distance between embeddings of the previous and current foveal fixation patches in a ResNet50 trained with SimSiam. Novelty is extracted at 4 main layers to measure visual features across the visual hierarchy. C) Correlation matrix of saccade amplitude and hierarchical novelty features from the human scalp EEG dataset. Novelty features are strongly correlated with each other, as well as saccade amplitude. D) Saccades and fixations are represented as pulses in time, and features associated with each as scaled pulses. The model finds a temporal response function (TRF, i.e. a linear impulse response) for each feature that best predicts the EEG. Modeling performance is measured with Pearson’s correlation r.

### Temporal response function encoding models

To determine how novelty modulates neural responses around eye movements, we used linear encoding models to predict neural activity at each electrode as output, using a time-lagged representation of eye-movement events and features as input (Holdgraf et al., 2017). The weights of these models are often called temporal response functions (TRFs), as the weights correspond to the brain’s responses that are linearly predictable from the input (Crosse et al., 2016). After these weights are learned, the model fit can be tested in a cross-validated manner to determine how well the model’s predictions of neural activity match the true neural activity using Pearson correlation. The use of cross-validation, consistent TRF amplitudes when adding features, and a consistent relationship between the zero-order correlation between novelty and EEG and novelty TRF weights all indicate our results are not due to suppression effects (Conger, 1974).

#### Banded ridge regression

Ridge regression is often used in encoding models because regularization helps prevent overfitting due to limited or noisy data. Many previous studies use the same regularization parameter for the entire combined feature space. However, it is not clear that this is the best solution, given that feature spaces with different units, different numbers of dimensions, different degrees of correlation, different degrees of sparsity, and/or different degrees of predictive power will require different regularization values to prevent overfitting (De Heer et al., 2017). Therefore, we use banded ridge regression to set a separate regularization value, λ, for each feature space (Nunez-Elizalde et al., 2019; Dupré La Tour et al., 2022). To do so, a cross-validation procedure is used, involving a search over each set of λ values for each feature space. This is done in a pre-specified order such that any shared variance will be attributed to the feature added first.

This is implemented by sweeping λ_1_ for the first feature, identifying λ_1_ that maximizes the average cross-validated correlation between the true and predicted neural responses, and freezing λ_1_ for the first feature. Then, the next feature is concatenated to the input, and now λ_2_ is swept for the second feature while holding λ_1_ constant to identify the λ_2_ that maximizes the prediction correlation. This process is repeated for each feature. This sequential approach assures us that any variance explained by an additional feature is not due to correlations with previously added features. We have used this same procedure in past work (Raghavan et al., 2023; Raghavan and Parra, 2025).

#### TRF modeling of feature hierarchy

All models were trained using the sklearn Ridge model solved via Cholesky decomposition and used cross-validation (CV) within participants to prevent overfitting (Pedregosa et al., 2011).

The movie datasets used leave-one-trial-out CV to avoid concatenation of discontinuous data. Trials are movie chapters in the human scalp dataset and movie clips in the human and NHP intracranial datasets. The image datasets used a 5-fold CV to ensure sufficient data was present in each fold.

The human scalp dataset used a time range of -0.2 to 0.8 seconds to capture the main components of each response, while limiting the extent to avoid unnecessary computational overhead due to the large size of the dataset. Fifteen regularization values, λs, were tested on a logarithmic scale from 10^-2^ to 10^5^. The human and NHP intracranial datasets used a time range of -0.5 to 1.0 seconds to capture the full BHA response of electrodes with a wide range of latencies. Ten regularization values, λs, were tested on a logarithmic scale from 10^0^ to 10^3^.

Regularization was applied by dividing the input features by their corresponding λ, as this is equivalent to multiplying the regularization term by λ (Nunez-Elizalde et al., 2019).

To measure the effect of each hierarchical novelty feature on neural responses invariant to the others, we used banded ridge regression with full and reduced models. First, we constructed the full model by adding features in the following order: saccade onset, fixation onset, saccade amplitude, and the combined novelty feature. We used a combined novelty feature with layer 1, 2, 3, and 4 so these features would have the same regularization. Then, we computed reduced models by replacing the combined novelty feature with a reduced novelty feature lacking the layer to be measured. For example, to measure the effect of layer 4 novelty, we measured the difference in correlation between the full model and the reduced model that uses layers 1, 2, and 3 novelty in the reduced novelty feature. TRF weights were analyzed from the full model.

#### Statistical testing of model predictions and TRF weights

Statistical tests performed to test whether novelty significantly modulated neural responses depended on the dataset. In the human scalp dataset, we first tested whether novelty was significantly encoded in neural activity using the hierarchical bootstrap of the encoding results, i.e., prediction improvement Δr, over participants and chapters (N_boot_ = 10^4^)(Saravanan et al., 2020). To confirm the TRF weights representing the novelty response showed significant differences from zero, we performed a non-parametric spatiotemporal cluster permutation test utilizing the adjacency matrix of the BioSemi64 montage and N=10^4^ permutations (Maris and Oostenveld, 2007). This revealed four clusters with permutation p<0.05 (SFig 2D), which include peaks at the noted time points (Fig 2E). In the human and NHP intracranial datasets, we tested whether novelty was significantly encoded at each electrode. To do so, we bootstrap the encoding results over trials and apply false discovery rate (FDR) correction to the resulting values (Benjamini and Hochberg, 1995; Saravanan et al., 2020). Only electrodes with p_boot_ < 0.05 with FDR correction are considered significant.

**Figure 2.**
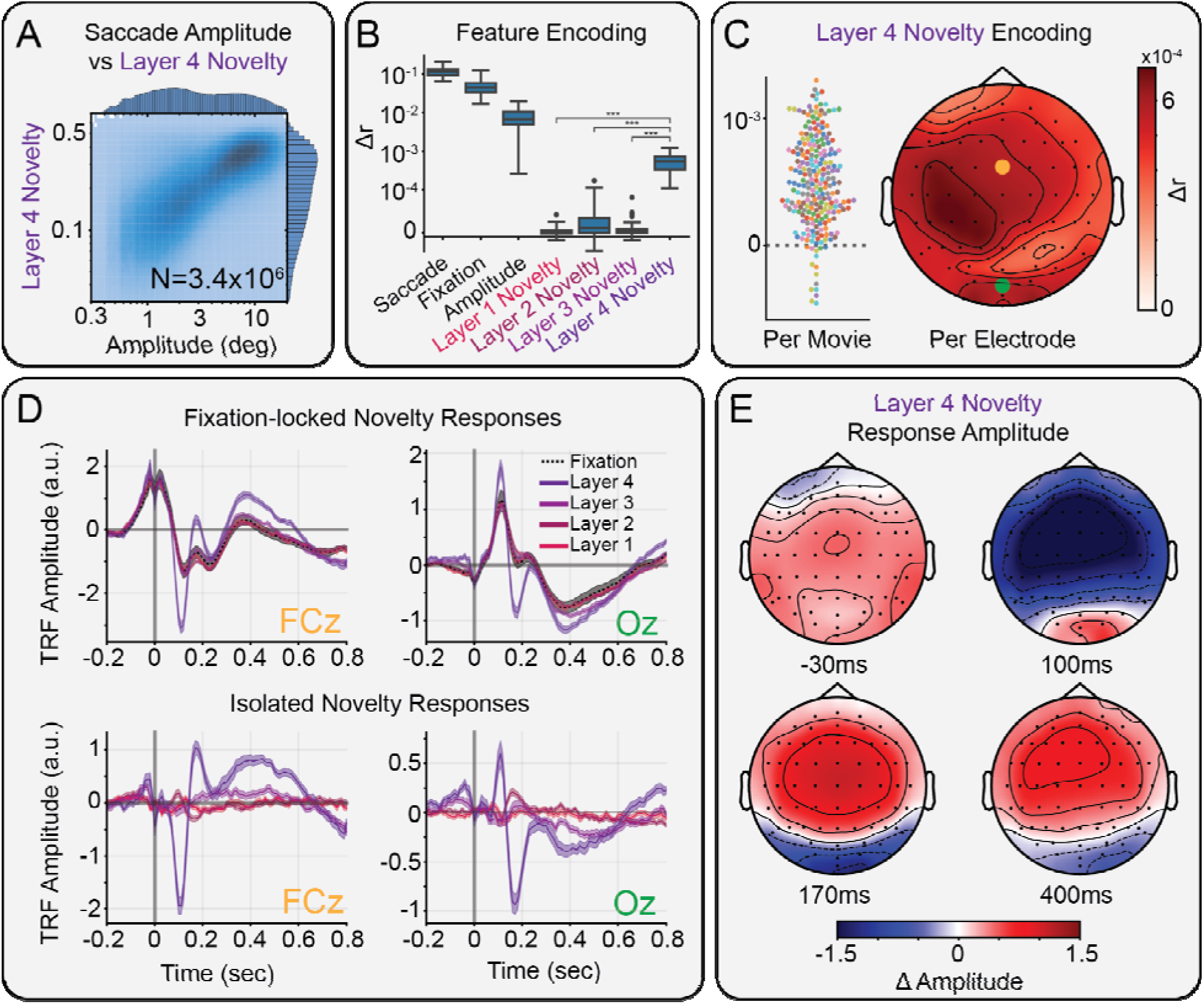
Modulation of scalp EEG by semantic novelty. A) Histogram of saccade amplitude vs layer 4 novelty, showing a correlation over N=3.4 million saccades from 231 movie viewings in 79 participants. B) Box plots over participants of the increase in correlation between predicted and actual EEG, when adding each feature to the encoding model. Layer 4 novelty captures significantly more variance than novelty computed from earlier layers (***: p<10^-3^). C) Increase in correlation from adding layer 4 novelty to the encoding model for each movie watched (left) and for each EEG electrode (right). Each dot is a viewing for a full-length film (90min - 120min). Color indicates the different movies (N=10). D) Average fixation-locked TRFs of electrodes FCz (left, yellow) and Oz (right, green). The additive effect of novelty (color) modulating the fixation response (dashed black; top). The isolated effect of novelty on fixation-locked responses (bottom). Different color lines from red to purple show layer 1 to layer 4 novelty responses. Only layer 4 novelty modulates neural activity. E) Topographic maps showing the modulation of components of fixation-locked responses by layer 4 novelty at -30, 100, 170, and 400ms.

#### Comparing saccades and fixations in human scalp EEG

Given that related work differs between the usage of saccade- and fixation-locked brain activity (Ries et al., 2018; Stankov et al., 2021; Nentwich et al., 2023; Madison et al., 2025), we compared approaches in the human scalp EEG dataset, time-locking features to either saccade or fixation, to assess how these predictors compare in their capacity to explain neural activity.

First, we trained encoding models using only saccade-onset and only fixation-onset predictors to see which better predicts neural activity. Next, we trained another pair of models, this time comprising onset and amplitude locked to saccades or fixations, and we analyzed how much the predictions of neural activity improved with amplitude. Specifically, we measured how much the Pearson correlation between true and predicted responses increased when adding amplitude beyond the onset model, i.e., the change in correlation between the onset-only model and the onset+amplitude model. We then compared those increases between the saccade- and fixation-locked models. We then trained another pair of models comprising onset, amplitude, and semantic novelty, locked to either saccades or fixations. As before, we measure how much predictions of neural activity improved with semantic novelty by comparing the onset+amplitude model and the onset+amplitude+novelty model for both saccade- and fixation-locked events.

### Brain plots

Scalp EEG topoplots were constructed using the MNE topomap visualization and BioSemi64 montage (Gramfort, 2013). Contour lines are shown along interpolated constant values for each scalp map. Human intracranial EEG brain plots were constructed using the electrode localization and plotting features in naplib-python (Mischler et al., 2023). Specifically, naplib can use Freesurfer-defined atlases to provide anatomical labels of electrodes based on FSAverage coordinates, as well as interpolate individual values onto the brain and summarize within atlas regions (Fischl, 2012). Here, we used the Desikan-Killiany atlas for displaying the percent electrodes predicted significantly and electrode latency, as this is a relatively coarse atlas showing broad effects (Desikan et al., 2006). The latency values for each region were obtained by taking the average latency of all electrodes labeled in that region. The Glasser atlas was used to display the modulation of individual electrodes over the brain, as this is more fine-grained and allows the identification of more relevant subregions (Glasser et al., 2016). The modulation values for each region were obtained by first interpolating the modulation value of each electrode onto the neighboring area using inverse distance weighting (max. distance 8mm) and then taking the average over all points in each region.

## Results

We analyzed scalp and intracranial EEG (iEEG) in humans and non-human primates (NHP) across three free-viewing datasets of movies and images (Fig 1A). These included human scalp EEG while watching full-length feature films (79 participants), human iEEG while viewing a series of ∼5-minute video clips and 6-second image presentations (31 participants), and NHP iEEG while viewing the same video clips and images (2 subjects). Most of these data (95.2%) have not been previously analyzed. Neural activity was measured via scalp potentials or broadband high-frequency amplitude (BHA).

To test the hypothesis that the brain responds to semantic novelty between fixations, we computed hierarchical novelty values from each 5° foveal fixation patch and analyzed how these differences modulate fixation-locked neural activity. We measured the modulation of neural activity using cross-validated linear encoding models (Fig 1D)(Crosse et al., 2016; Holdgraf et al., 2017; Ehinger and Dimigen, 2019). This method learns a set of weights that linearly predict neural activity from a time-lagged version of the input. The model weights correspond to the impulse response of each feature, also referred to as the temporal response function (TRF). If adding a feature to the model significantly improves predictions of neural activity, i.e. there is a significant increase in Pearson correlation Δr, then we conclude the brain represents or “encodes” this feature.

To compute hierarchical representations of each fixation patch, we passed the patch through a ResNet50 trained with SimSiam (Chen and He, 2021) and extracted the embeddings at four main layers of the network (Fig 1B). We selected SimSiam as it is trained entirely with prediction and does not require any labels. This resembles the self-supervised prediction aspects we postulate for primate visual systems. Using these four layers provides a range of representations of each patch derived directly from a model trained with unsupervised learning, i.e., to minimize distances between representations (embeddings) of different patches from the same image. Novelty for each layer is computed as the cosine distance between the embeddings of successive foveal fixation patches at that layer.

In the scalp EEG dataset, we find strong correlations between saccade amplitude and novelty from different layers, with Layer 4 novelty being strongest (Fig 1C). Indeed, in all three datasets, we find that Layer 4 novelty is strongly correlated with saccade amplitude (Fig 2A, 3A, 4A). This is unsurprising, as more distant locations are likely to capture visually distinct elements, and even more likely to capture semantically distinct elements in natural scenes. The strong correlation between novelty computed from different layers indicates that these features are capturing relevant low- and mid-level visual differences that are closely related to the semantic differences observed at Layer 4. We find that a range of low- and mid-level visual features from each fixation patch, as well as saliency and optic flow, generally do not correlate with our novelty measures (Fig S1)(Dorr et al., 2010; Itti, 2005; Itti and Koch, 2000). The analysis of fixation-related responses therefore only included saccade amplitude and novelty features, as they are strongly correlated and related to the full visual hierarchy.

**Figure 3.**
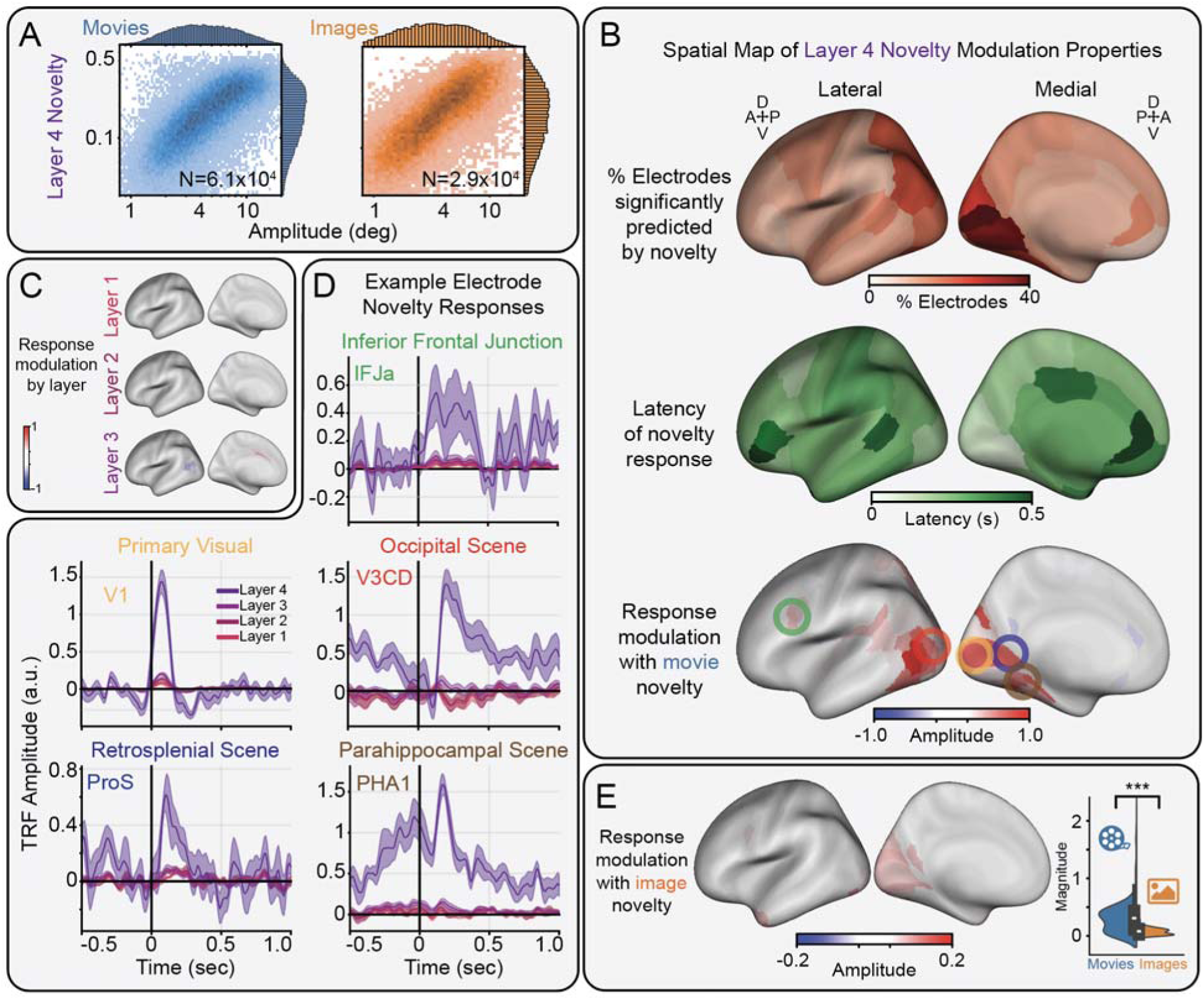
Modulation of human intracranial EEG by semantic novelty. A) Histogram of saccade amplitude vs layer 4 novelty, showing a correlation over N=60,550 and N=28,797 saccades in the movie-(left) and image-viewing (right) datasets, respectively. B) Spatial distribution of layer 4 novelty modulation properties, including the percent of electrodes significantly encoding layer 4 novelty (top), the average latency of sites responding to layer 4 novelty (center), and the maximum modulation for sites responsive to layer 4 novelty during movie-viewing (bottom). Green, yellow, red, blue, and brown circles correspond to the regions shown in panel D. C) Maximum modulation for sites responsive to layers 1, 2, and 3 novelty during movie-viewing. D) Time course of novelty modulated fixation responses during movie viewing in selected brain regions, including the inferior frontal junction (IFJ), primary visual cortex (V1), as well as the occipital (V3CD), retrosplenial (ProS), and parahippocampal (PHA1) scene areas (mean ± s.e.). E) Spatial distribution of the maximum modulation for sites responsive to layer 4 novelty during image-viewing (left). Modulation magnitude of sites responsive to layer 4 novelty during movie-viewing is significantly greater than during image-viewing (***: p<0.001; right).

**Figure 4.**
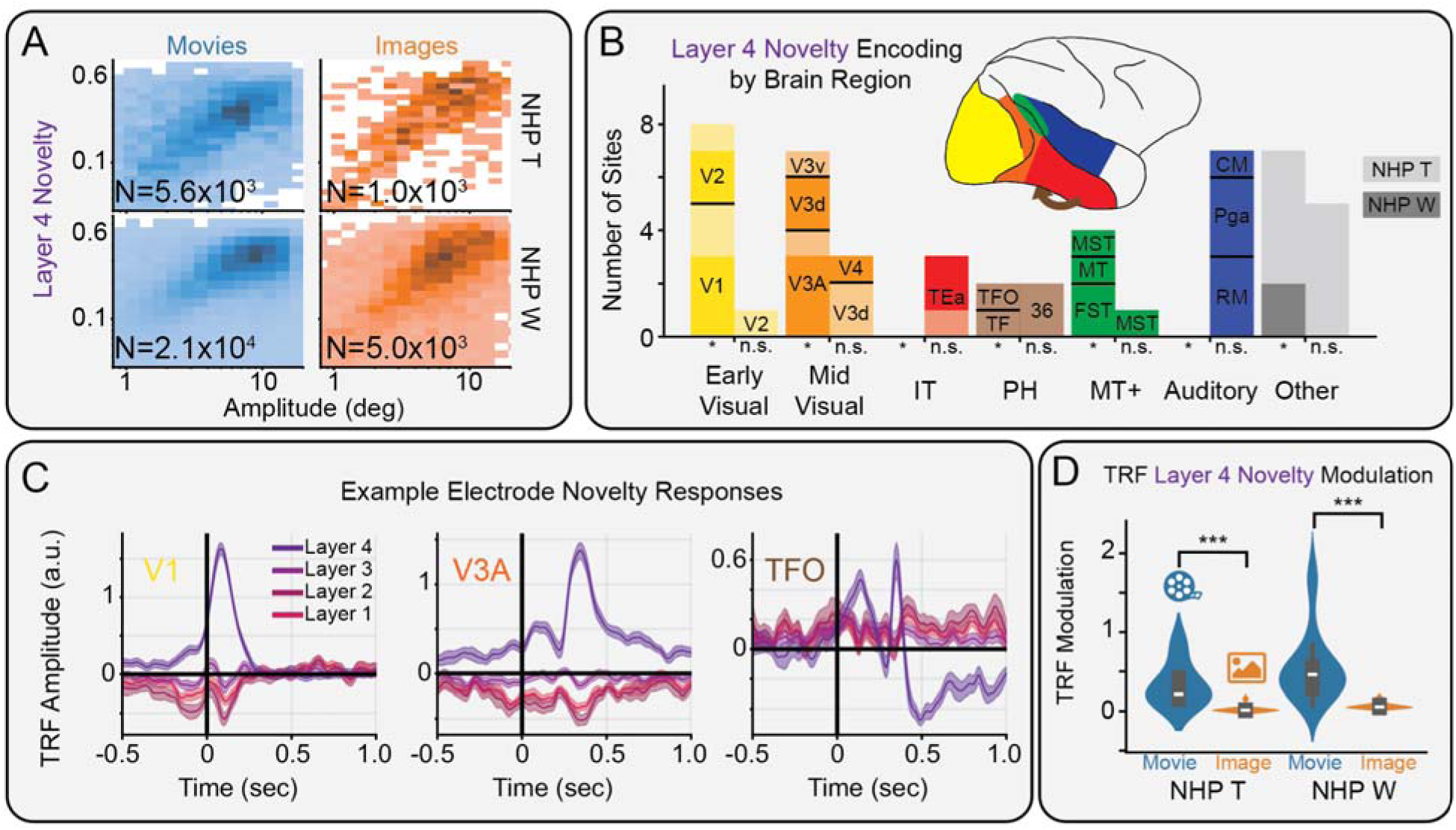
Modulation of non-human primate intracranial EEG by semantic novelty. A) Histogram of saccade amplitude and layer 4 novelty, during movie (left) and image viewing (right) for NHP T (top) and NHP W (bottom). B) Bar plot showing where electrodes were located, including those with (left bars) and without (right bars) significant (*: p_boot_ < 0.05, FDR corrected) modulation by novelty in NHP T (light shading) and NHP W (dark shading). C) Time course of novelty modulation in selected areas along the visual hierarchy, including V1, V3A, and TFO. D) Modulations of novelty TRFs are significantly (***: p<0.001) stronger during movie-vs image-viewing.

### Only semantic novelty modulates fixation-locked human scalp EEG

We measured saccade- and fixation-related responses using TRFs in human scalp EEG. Saccades and fixations were coded as impulses to separate their neural contributions (Fig 1E). Saccade amplitude was aligned to saccade onset, while all novelty features were aligned to fixation onset. This alignment best explained the variance in the EEG (Fig S2A). After accounting for saccade amplitude with non-linear responses (Dimigen and Ehinger, 2021), novelty significantly modulated fixation-locked but not saccade-locked responses (Fig S2C) confirming that novelty modulation is tied to fixational processing. As expected, most variance is explained by the saccade and fixation regressors, with additional variance explained by the features that modulate these fixed responses (quantified as Δr in Fig. 2B; for total variance explained see Fig S2E). Notably, layer 4 novelty captures significantly more variance than layer 1 novelty (t(78)=17.4, p=4.7×10^-29^), layer 2 novelty (t(78)=17.1, p=1.4×10^-28^), and layer 3 novelty (t(78)=17.4, p=3.9×10^-29^; paired t-test, Holm-Bonferroni corrected).

We measured how much layer 4 novelty modulates fixation-locked responses. Across the 231 movie viewings (Fig 2C, left) and 64 scalp electrodes (Fig 2C, right), we found significant variance explained, particularly in frontal and occipital areas. Indeed, given the large number of saccades available, all 64 scalp electrodes showed significant modulation (all Δr>3.1×10-4, p_boot_<0.05, FDR-corrected). The spatial distribution of variance explained on the scalp (Fig. 2C) rules out eye movement artifacts as the source of this modulation. Inspecting the TRFs for individual electrodes reveals how novelty modulates particular components of the underlying fixation-related responses, with similar spatial and temporal profiles to the underlying response (Fig 2D).

To characterize the spatiotemporal effects of novelty, we plotted the spatial distribution of the novelty TRFs at local extrema (Fig 2E). These extrema were identified in clusters with permutation p<0.01 by a non-parametric spatiotemporal cluster permutation test (Fig S2D)(Maris and Oostenveld, 2007). A fronto-central modulation precedes fixation onset by 30ms. At 100ms after fixation onset, we find the largest modulation of frontal areas. At 170 and 400ms after fixation onset, we find fronto-occipital dipoles, including occipital negativities. These modulations match the sign of the baseline fixation-related potentials and thus represent an overall enhancement, rather than suppression, of the underlying neural response.

### Semantic novelty modulates human intracranial EEG in scene-selective regions along the ventromedial visual stream

We analyzed eye movement responses in the BHA of human iEEG in N=6328 electrodes with coverage over the entire brain (Fig S3). The same extraction and modeling approaches were used to quantify hierarchical novelty features from fixation patches and measure the isolated effects of novelty at different levels of abstraction (Fig 3A). Electrodes containing saccadic spike artifacts due to extraocular muscle activity (Jerbi et al., 2009) were identified via hierarchical clustering of the correlation matrix of saccade TRFs and removed from further analysis (Fig S3B)(Yeo et al., 2011). We found that 12.4% of the remaining sites (708/5693) significantly encoded layer 4 novelty, primarily in occipital areas (p_boot_<0.05, FDR-corrected; Fig 3B, top).

The peak latency of novelty modulations across the cortex were quantified as the time of the absolute maximum of the signal, i.e., the highest peak or lowest trough. These peak latencies were interpolated onto each region of the Desikan-Killiany atlas (Fig 3B, center). This reveals two trends: an increasing latency along the ventro-medial visual stream from posterior to anterior areas, and an early frontal response in the caudal middle frontal area, which lies between the inferior frontal junction (IFJ) and the frontal eye fields, regions responsible for controlling visual attention (Bedini & Baldauf, 2021).

We analyzed the modulations of electrodes significantly encoding novelty. On average, these sites favored enhancement (57%) rather than suppression (43%) of the BHA activity. The amplitude of the layer 4 novelty modulation, defined as the highest peak or lowest trough, was then interpolated onto the HCP-MMP atlas of the brain (Fig 3B, bottom). This revealed that enhancements dominate the strongest modulations in IFJ, as well as along the ventromedial visual stream, particularly in scene-selective visual regions. Brain plots of the modulation amplitude for novelty computed at layers 1, 2, and 3 reveal limited modulations due to these measures of novelty (Fig 3C).

We plotted the layer 4 novelty TRF of exemplar electrodes in each of these regions (Fig 3D). We note strong enhancements in the IFJ, a region responsible for guiding feature-based, rather than spatial, visual attention. Strong enhancements are also observed in primary visual cortex V1 and each of the scene-selective visual areas, including the occipital, retrosplenial, and parahippocampal scene areas, corresponding to electrodes in HCP-MMP atlas regions V3CD, ProS, and PHA1, respectively. These responses represent an enhancement of the underlying fixational BHA responses (Fig S3C).

An identical analysis was performed on the human iEEG image-viewing dataset (N=8522 electrodes) to determine if novelty modulation persists during image-viewing. We found that 11.5% of sites without saccade artifacts (894/7803) significantly encoded layer 4 novelty. These sites had a spatially similar but significantly (t(1,600)=23.2, p=9.8×10-103; unpaired t-test) decreased magnitude of layer 4 novelty modulation across significantly modulated electrodes in each dataset (Fig 3E). Overall, these results suggest semantic novelty enhances frontal areas responsible for visual attention, primary visual cortex, as well as scene-specific areas along the ventromedial visual stream during movie-watching more than image-viewing.

### Semantic novelty modulates NHP iEEG along the ventral visual stream during movie-watching more than image-viewing

Finally, we analyzed BHA responses to eye movements during movie-watching in iEEG recorded from two NHPs with 50 implanted sEEG contacts broadly covering temporal and occipital cortical regions. As before, we measured the effect of hierarchical novelty features on neural activity beyond saccade amplitude (Fig 4A). We find that 28/50 (NHP T: 12/23, NHP W: 16/27) sites are significantly modulated by layer 4 novelty, while these modulations are weaker for layers 1, 2, and 3 (Fig 4B). Modulations of sites in visual regions favored enhancement (NHP T: 5/7, NHP W: 11/14) over suppression of BHA activity. This includes early, middle, and late responses along the expanded ventral visual pathway in V1, V3A, and TFO, respectively (Fig 4C)(Kravitz et al., 2013). Notably, areas V3A and TFO are both thought to be scene-selective in macaques (Kornblith et al., 2013).

To examine the influence of a dynamic vs static stimulus, we compared the layer 4 novelty modulation of neural responses during movie viewing with those during image viewing. We found that novelty modulation was significantly (NHP T: t(11)=5.56, p=1.7×10^-4^; NHP W: t(15)=5.59, p=5.1×10^-5^; unpaired t-test, Holm-Bonferroni corrected) reduced across novelty-modulated sites during the presentation of static images compared to movie clips, consistent with our results in human iEEG (Fig 4D). Overall, these results suggest that semantic novelty enhances activity along scene-specific regions in the NHP ventral visual stream following fixation onset during natural viewing of movies more than images.

## Discussion

Our study demonstrates that semantic novelty enhances scalp EEG fixation-locked potentials and broadly enhances human and NHP iEEG BHA activity, particularly in scene-selective visual regions. This spatial profile persists between image and movie viewing. Frontal areas, including the inferior frontal junction, show early modulation while occipital modulation occurs both early and late. These findings indicate the visual system extracts foveal semantic information and integrates it between fixations to build a representation of the scene.

### Similarities to known EEG evoked response components

Fixed-gaze ERPs often parallel free-viewing fixation-related responses (Kamienkowski et al., 2012). Our data reveal modulations mirroring established novelty components: The mismatch negativity (MMN), a response to stimulus change that occurs without attention (Pazo-Alvarez et al., 2003); the P3a, a fronto-central response reflecting orienting to novel stimuli (Friedman et al., 2001); and the N400, a response to semantic incongruity in a range of domains (Kutas and Federmeier, 2011). We observe a P3a-like fronto-central positivity *before* fixation onset, consistent with the orienting process anticipating novelty before fixation. Next, we see fronto-occipital responses, including occipital negativities around 170ms and 400ms. While the frontal positivities appear to dominate the scalp maps, the occipital negativities observed more directly relate to those expected from novelty responses from the visual cortex, like the visual MMN and visual N400 (Barrett and Rugg, 1990; Astikainen et al., 2013).

### Early frontal modulation, early and late occipital modulation

We find modulations in frontal areas near fixation onset. Early frontal modulation suggests novel semantics are processed in the periphery, from pre-saccadic attention shifts, or possibly anticipated from context, aligning with work showing novelty signals during the *preceding* fixation (Coco et al., 2020). This suggests the brain is not only predicting the upcoming input, but also how novel the upcoming input might be. Conversely, late modulation in early visual areas suggests top-down feedback, where foveal areas respond to peripheral inputs after a delay (Stewart et al., 2020). The timing of this effect is consistent with the integration of peripheral signals with subsequent foveal input (Ramezani et al., 2019). Modulation of early areas with semantic novelty is consistent with fMRI studies showing that object category information and high-level prediction error are fed back to foveal retinotopic visual cortex and early visual cortex, respectively (Williams et al., 2008; Richter et al., 2024).

### Semantic novelty modulation in scene-selective regions indicates the function of integration

Our results revealed strong modulation with novelty in the parahippocampal, retrosplenial, and occipital scene areas in humans (Epstein and Baker, 2019) and areas V3A and TFO in NHPs, also associated with scene processing (Kornblith et al., 2013). This common modulation suggests these streams are conserved across species (Vinken et al., 2025), potentially including the ventral novelty stream (Weierich et al., 2010; Kafkas and Montaldi, 2018). The activation of scene-selective regions indicates that novel fixations help build scene representations. This agrees with the importance of saccade targets for transsaccadic integration in natural scenes (Henderson and Hollingworth, 2003) and the role of oculomotor cues for integration in scene-selective regions (Golomb et al., 2011). Occipital and parahippocampal regions also respond to eye movements during movie recall, suggesting that recall involves the neural reinstatement of both visual content and the dynamics of its integration (Nau et al., 2025).

### Stronger responses to movies vs images

Novelty responses in movies were stronger than in images throughout the cortex, aligning with existing studies showing stronger responses to dynamic vs static stimuli for faces (Pitcher et al., 2011), bodies and objects (Küçük et al., 2024), and scenes (Korkmaz Hacialihafiz and Bartels, 2015). This typically occurs in brain areas sensitive to motion, such as integrated motion in area MT/V5 (Born and Bradley, 2005), self-motion and optic flow in MST (Wild and Treue, 2021), and “biological motion” in STS (Blake and Shiffrar, 2007) but rarely in the ventral visual stream (Pitcher et al., 2019).

While motion could contribute, past work shows motion has a comparatively weak effect (Nentwich et al., 2023). In contrast, cuts elicit large responses, likely due to sudden changes in the scene. We control for this by excluding saccades crossing cuts. Instead we propose alternative explanations: First, movies increase engagement and attention to the stimulus, increasing evoked activity. Second, our novelty measure based on static patches underestimates novelty due to dynamics in the foveal input, leading to an overestimate of modulation during movies. Third, our local estimate of novelty may correlate with global novelty, and thus, capture broader responses due to changes in the whole scene. However, the novelty response observed here cannot be uniquely attributed to global changes as we observed the same, albeit weaker effects during static image-viewing. Future work could explore how local (foveal) versus global (scene) novelty modulate neural activity.

### Possible role of enhancing responses with novelty

We found that novelty primarily enhanced the underlying fixation responses, showing similar spatial and temporal profiles. While probabilistic population coding suggests expected stimuli should evoke stronger responses (Ma et al., 2006), most temporal prediction models argue for weaker responses (Summerfield and de Lange, 2014). In V1, expected stimuli cause weaker fMRI responses but can be more readily decoded, aligned with a “sharper” representation (Kok et al., 2012). The theory of “predictive coding” argues that “error” units compare top-down prediction with bottom-up input, responding to mismatches (de Lange et al., 2018). This theory argues that the error signal propagates forward and serves as the primary “encoder” of the sensory input (Millidge et al., 2022). However, the biological realism of this error-coding mechanism is not universally accepted (Walsh et al., 2020; Solomon et al., 2021; Westerberg et al., 2024). More widely accepted is the notion that prediction errors are relevant for learning (Roelfsema and Holtmaat, 2018).

### Error signal as a potential tuning signal

This work was motivated by recent progress in machine learning showing that vision models can be trained without labels, but simply by predicting representations of one image from the next. The concept has been incorporated in a “joint embedding predictive architecture” (JEPA) (LeCun, 2022), which is based on predicting representations of the upcoming input using the current input and action. While SimSiam uses prediction (Chen and He, 2021), JEPA makes the temporal prediction explicit. Recent work has shown that the training objective of predicting the embedding of the next input produces representations more aligned with the ventral visual stream (Thorat et al., 2026). In this framework, the error in predicting the upcoming embedding is the training signal to tune the network. If a similar mechanism is at play in the brain, we expect ubiquitous novelty responses, which indeed we found. The observed early frontal modulation even suggests this signal may be anticipated. Vision remains adaptive throughout life (Webster, 2015). Indeed, evidence suggests prediction between peripheral and foveal vision serves to calibrate size perception (Valsecchi and Gegenfurtner, 2016). More generally, we speculate that semantic novelty, which can be considered a prediction error, may serve to tune the ventral visual system engaged in scene representation.

### Limitations

Our measure of novelty did not account for peripheral vision, which influences foveation targets (Stewart et al., 2020). Psychophysics research shows integration of the current peripheral with upcoming foveal vision (Deubel and Schneider, 1996). Neurons responsive to foveal vision respond, shortly before a saccade, to the to-be-fixated stimulus already visible in peripheral vision (Duhamel et al., 1992; Neupane et al., 2020). Evidence for semantic processing of peripheral vision also comes from scalp recordings with neural responses to the target *prior* to the upcoming fixation (Ehinger et al., 2015; Buonocore et al., 2020; Stankov et al., 2021). Future studies should explore measures of novelty that incorporate peripheral vision. Similarly, this measure of novelty only measures integration across consecutive fixations, rather than across a whole sequence. Future work can use recurrence in a measure of novelty to determine if novelty integrated over a sequence of fixations is a better predictor of fixation-related potentials during scene viewing (Thorat et al., 2025).

Finally, solving for the regularization separately for each feature and holding it constant, allows us to isolate contributions of each feature. However, we do not re-optimize lambdas when adding features, meaning the final model may be non-optimal. Therefore, our reported contributions are more conservative than expected from a joint optimization.

## Conclusion

Our findings provide compelling evidence that the visual system computes the semantic novelty of fixations. This novelty is reflected in the modulation of fixation-related responses across multiple brain regions and in both humans and non-human primates. By controlling for saccade amplitude and hierarchical novelty features, we isolate a purely semantic novelty that likely serves to tune the ventral visual system, including scene-specific areas. This is consistent with our postulate that subsequent fixations integrate semantic information to build a representation of the scene. These results challenge traditional models of vision that treat each fixation as an independent snapshot and highlight the importance of considering the temporal dynamics of eye movements and the predictive nature of visual processing.

## Acknowledgements

We would like to thank Alexandria Hoang and Angel Him Lee who contributed to the human scalp EEG data collection; Ashesh Mehta, Noah Markowitz, Elizabeth Espinal, Gelana Tostaeva, Sabina Gherman and Jose Herrero, who contributed to the human iEEG collection; Brent Butler, Kurt Masiello, and Debbie Ross who contributed to the NHP iEEG data; and Charles E. Schroeder who contributed to funding acquisition. This work was supported in part by grants ARO WF911-NF-24-1-0031, NIH P50 MH109429, and NSF DRL-2201835.

## Supplemental Figures

**Figure S1.**
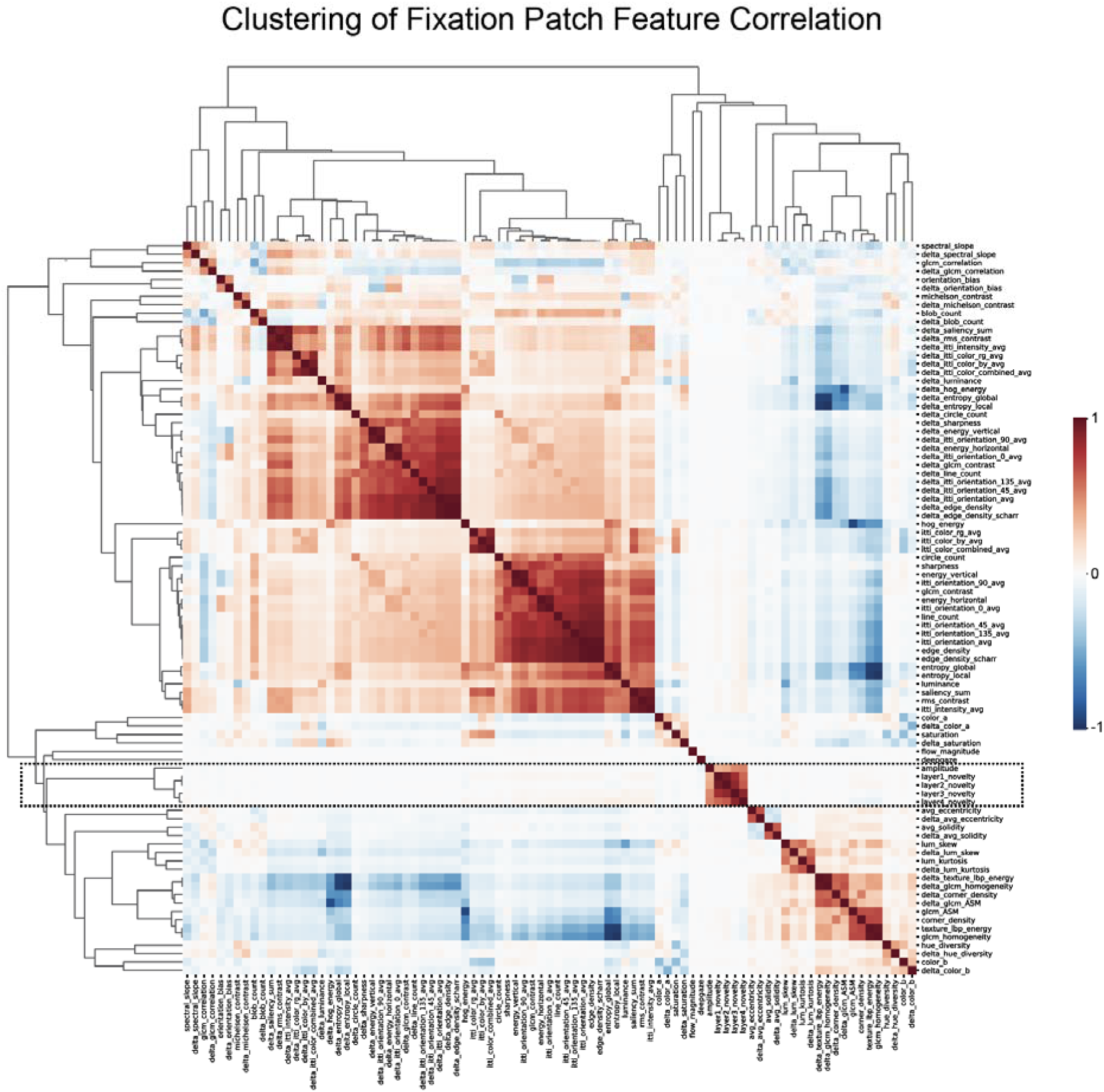
Correlation of hierarchical novelty with other visual features. Hierarchical novelty features are only strongly correlated with saccade amplitude (black dashed box).

**Figure S2.**
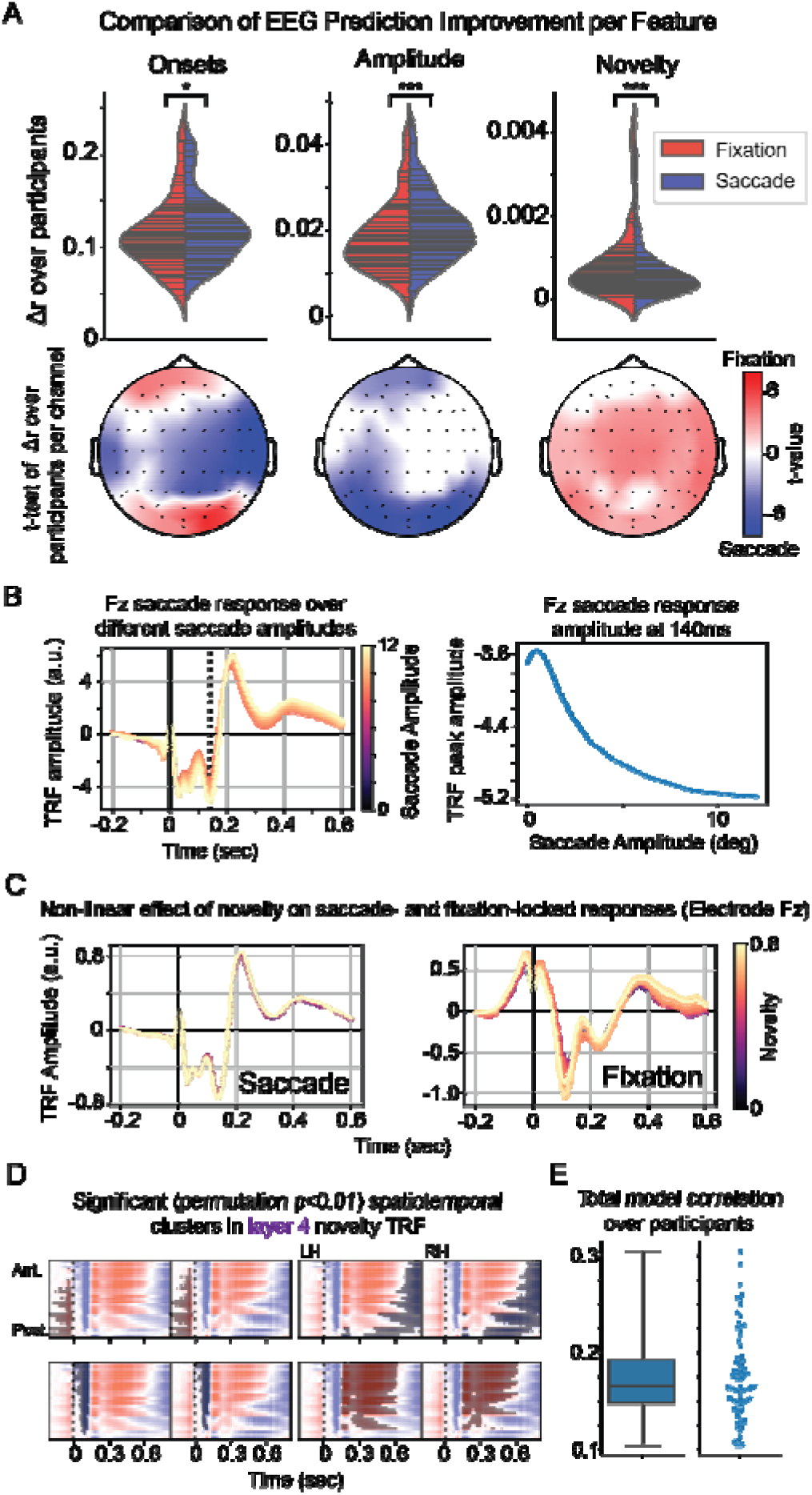
Modulation of scalp EEG potentials. A) Left: Saccade onsets capture slightly more variance than fixation onsets (*: p<0.05), and different electrodes are better explained by saccades vs fixations. Center: Amplitude locked to saccade onset captures more variance than amplitude locked to fixation onset (***: p<0.001), and this effect is consistent across the scalp. Right: Novelty locked to fixation onset captures more variance than novelty locked to saccade onset, with amplitude regressed out (***: p<0.001), and this effect is consistent across the scalp. B) Saccade amplitude responses modeled with B-splines display a non-linear response with saccade amplitude. C) Novelty only modulates fixation-locked responses with saccade amplitude regressed out. D) Significant clusters in layer 4 novelty TRF (permutation p<0.01). E) Total TRF model correlation with all features added.

**Figure S3.**
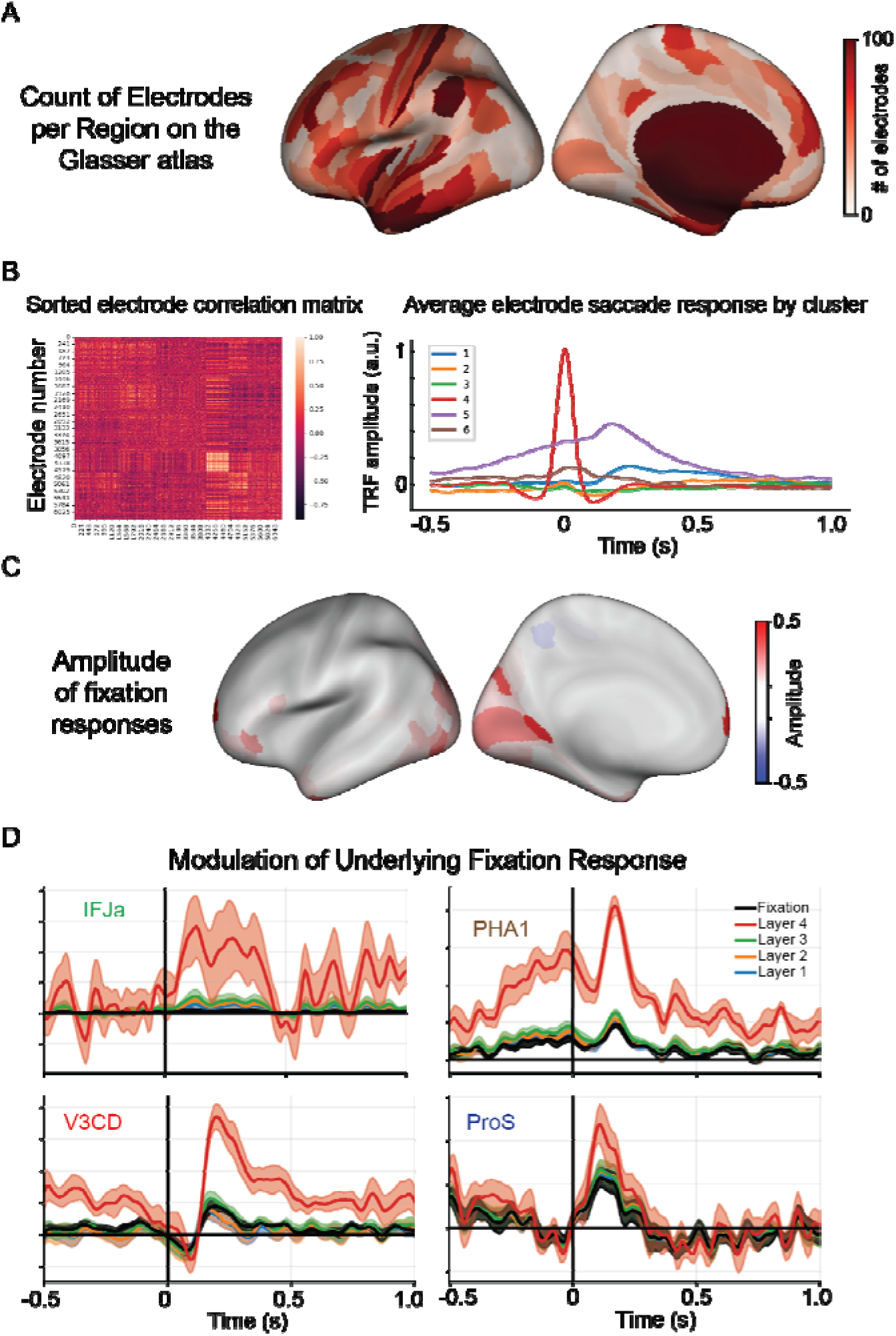
Human intracranial EEG coverage, electrode response clusters, and novelty enhancement vs suppression. A) Brain plot showing the total number of electrodes measured from each region of the Glasser atlas in the movie-watching dataset. B) Correlation matrix of electrode saccade response, sorted by clusters (left). Electrode saccade response, averaged within each cluster. Cluster 4 was identified as saccadic spike artifacts and removed from analysis. C) Amplitude of fixation responses interpolated onto the brain. This spatial layout is consistent with the novelty modulation; however, novelty shows particularly strong modulations in scene-specific regions not otherwise present for fixation responses. D) Novelty responses added to the underlying fixation BHA response in frontal and scene-specific regions.

## References

Astikainen P, Cong F, Ristaniemi T, Hietanen JK (2013) Event-related potentials to unattended changes in facial expressions: detection of regularity violations or encoding of emotions? Front Hum Neurosci 7 Available at: http://journal.frontiersin.org/article/10.3389/fnhum.2013.00557/abstract [Accessed February 4, 2026].

Barczak A, Haegens S, Ross DA, McGinnis T, Lakatos P, Schroeder CE (2019) Dynamic Modulation of Cortical Excitability during Visual Active Sensing. Cell Rep 27:3447–3459.e3.

Barrett SE, Rugg MD (1990) Event-related potentials and the semantic matching of pictures. Brain Cogn 14:201–212.

Bartlett AM, Ovaysikia S, Logothetis NK, Hoffman KL (2011) Saccades during Object Viewing Modulate Oscillatory Phase in the Superior Temporal Sulcus. J Neurosci 31:18423–18432.

Benjamini Y, Hochberg Y (1995) Controlling the False Discovery Rate: A Practical and Powerful Approach to Multiple Testing. J R Stat Soc Ser B Methodol 57:289–300.

Blake R, Shiffrar M (2007) Perception of Human Motion. Annu Rev Psychol 58:47–73.

Born RT, Bradley DC (2005) STRUCTURE AND FUNCTION OF VISUAL AREA MT. Annu Rev Neurosci 28:157–189.

Buonocore A, Dimigen O, Melcher D (2020) Post-Saccadic Face Processing Is Modulated by Pre-Saccadic Preview: Evidence from Fixation-Related Potentials. J Neurosci 40:2305–2313.

Chen X, He K (2021) Exploring Simple Siamese Representation Learning. In: 2021 IEEE/CVF Conference on Computer Vision and Pattern Recognition (CVPR), pp 15745–15753. Nashville, TN, USA: IEEE. Available at: https://ieeexplore.ieee.org/document/9578004/ [Accessed July 30, 2025].

Coco MI, Nuthmann A, Dimigen O (2020) Fixation-related Brain Potentials during Semantic Integration of Object–Scene Information. J Cogn Neurosci 32:571–589.

Conger AJ (1974) A Revised Definition for Suppressor Variables: a Guide To Their Identification and Interpretation. Educ Psychol Meas 34:35–46.

Cox RW (1996) AFNI: software for analysis and visualization of functional magnetic resonance neuroimages. Comput Biomed Res Int J 29:162–173.

Crosse MJ, Di Liberto GM, Bednar A, Lalor EC (2016) The Multivariate Temporal Response Function (mTRF) Toolbox: A MATLAB Toolbox for Relating Neural Signals to Continuous Stimuli. Front Hum Neurosci 10 Available at: http://journal.frontiersin.org/article/10.3389/fnhum.2016.00604/full [Accessed June 12, 2025].

De Heer WA, Huth AG, Griffiths TL, Gallant JL, Theunissen FE (2017) The Hierarchical Cortical Organization of Human Speech Processing. J Neurosci 37:6539–6557.

de Lange FP, Heilbron M, Kok P (2018) How Do Expectations Shape Perception? Trends Cogn Sci 22:764–779.

Degno F, Liversedge SP (2020) Eye Movements and Fixation-Related Potentials in Reading: A Review. Vision 4:11.

Desikan RS, Ségonne F, Fischl B, Quinn BT, Dickerson BC, Blacker D, Buckner RL, Dale AM, Maguire RP, Hyman BT, Albert MS, Killiany RJ (2006) An automated labeling system for subdividing the human cerebral cortex on MRI scans into gyral based regions of interest. NeuroImage 31:968–980.

Deubel H, Schneider WX (1996) Saccade target selection and object recognition: Evidence for a common attentional mechanism. Vision Res 36:1827–1837.

Dimigen O, Ehinger BV (2021) Regression-based analysis of combined EEG and eye-tracking data: Theory and applications. J Vis 21:3.

Dorr M, Martinetz T, Gegenfurtner KR, Barth E (2010) Variability of eye movements when viewing dynamic natural scenes. J Vis 10:28–28.

Duhamel J-R, Colby CL, Goldberg ME (1992) The Updating of the Representation of Visual Space in Parietal Cortex by Intended Eye Movements. Science 255:90–92.

Dupré La Tour T, Eickenberg M, Nunez-Elizalde AO, Gallant JL (2022) Feature-space selection with banded ridge regression. NeuroImage 264:119728.

Ehinger BV, Dimigen O (2019) Unfold: an integrated toolbox for overlap correction, non-linear modeling, and regression-based EEG analysis. PeerJ 7:e7838.

Ehinger BV, König P, Ossandón JP (2015) Predictions of Visual Content across Eye Movements and Their Modulation by Inferred Information. J Neurosci 35:7403–7413.

Eisenberg ML, Zacks JM (2016) Ambient and focal visual processing of naturalistic activity. J Vis 16:5.

Epstein RA, Baker CI (2019) Scene perception in the human brain. Annu Rev Vis Sci 5:373–397.

Fischl B (2012) FreeSurfer. NeuroImage 62:774–781.

Friedman D, Cycowicz YM, Gaeta H (2001) The novelty P3: an event-related brain potential (ERP) sign of the brain’s evaluation of novelty. Neurosci Biobehav Rev 25:355– 373.

Glasser MF, Coalson TS, Robinson EC, Hacker CD, Harwell J, Yacoub E, Ugurbil K, Andersson J, Beckmann CF, Jenkinson M, Smith SM, Van Essen DC (2016) A multi-modal parcellation of human cerebral cortex. Nature 536:171–178.

Golomb JD, Albrecht AR, Park S, Chun MM (2011) Eye Movements Help Link Different Views in Scene-Selective Cortex. Cereb Cortex 21:2094–2102.

Gramfort A (2013) MEG and EEG data analysis with MNE-Python. Front Neurosci 7 Available at: http://journal.frontiersin.org/article/10.3389/fnins.2013.00267/abstract [Accessed February 18, 2025].

Groppe DM, Bickel S, Dykstra AR, Wang X, Mégevand P, Mercier MR, Lado FA, Mehta AD, Honey CJ (2017) iELVis: An open source MATLAB toolbox for localizing and visualizing human intracranial electrode data. J Neurosci Methods 281:40–48.

Hartig R, Glen D, Jung B, Logothetis NK, Paxinos G, Garza-Villarreal EA, Messinger A, Evrard HC (2021) The Subcortical Atlas of the Rhesus Macaque (SARM) for neuroimaging. NeuroImage 235:117996.

Hayes TR, Henderson JM (2021) Looking for Semantic Similarity: What a Vector-Space Model of Semantics Can Tell Us About Attention in Real-World Scenes. Psychol Sci 32:1262–1270.

Henderson JM, Hayes TR (2017) Meaning-based guidance of attention in scenes as revealed by meaning maps. Nat Hum Behav 1:743–747.

Henderson JM, Hollingworth A (2003) Eye movements and visual memory: Detecting changes to saccade targets in scenes. Percept Psychophys 65:58–71.

Holdgraf CR, Rieger JW, Micheli C, Martin S, Knight RT, Theunissen FE (2017) Encoding and Decoding Models in Cognitive Electrophysiology. Front Syst Neurosci 11:61.

Irwin DE, Gordon RD (1998) Eye Movements, Attention and Trans-saccadic Memory. Vis Cogn 5:127–155.

Itti L (2005) Quantifying the contribution of low-level saliency to human eye movements in dynamic scenes. Vis Cogn 12:1093–1123.

Itti L, Koch C (2000) A saliency-based search mechanism for overt and covert shifts of visual attention. Vision Res 40:1489–1506.

Jerbi K, Freyermuth S, Dalal S, Kahane P, Bertrand O, Berthoz A, Lachaux J-P (2009) Saccade Related Gamma-Band Activity in Intracerebral EEG: Dissociating Neural from Ocular Muscle Activity. Brain Topogr 22:18–23.

Jung B, Taylor PA, Seidlitz J, Sponheim C, Perkins P, Ungerleider LG, Glen D, Messinger A (2021) A comprehensive macaque fMRI pipeline and hierarchical atlas. NeuroImage 235:117997.

Kafkas A, Montaldi D (2018) How do memory systems detect and respond to novelty? Neurosci Lett 680:60–68.

Kamienkowski JE, Ison MJ, Quiroga RQ, Sigman M (2012) Fixation-related potentials in visual search: A combined EEG and eye tracking study. J Vis 12:4.

Kämmer L, Kroell LM, Knapen T, Rolfs M, Hebart MN (2026) Feedback of peripheral saccade targets to early foveal cortex. eLife 14:RP107053.

Kazai K, Yagi A (1999) Integrated effect of stimulation at fixation points on EFRP (eye-fixation related brain potentials). Int J Psychophysiol 32:193–203.

Kok P, Jehee JFM, de Lange FP (2012) Less Is More: Expectation Sharpens Representations in the Primary Visual Cortex. Neuron 75:265–270.

Korkmaz Hacialihafiz D, Bartels A (2015) Motion responses in scene-selective regions. NeuroImage 118:438–444.

Kornblith S, Cheng X, Ohayon S, Tsao DY (2013) A Network for Scene Processing in the Macaque Temporal Lobe. Neuron 79:766–781.

Kroell LM, Rolfs M (2026) Predictive Foveal Processing in Active Vision. Annu Rev Vis Sci 12.

Küçük E, Foxwell M, Kaiser D, Pitcher D (2024) Moving and Static Faces, Bodies, Objects, and Scenes Are Differentially Represented across the Three Visual Pathways. J Cogn Neurosci 36:2639–2651.

Kümmerer M, Bethge M (2023) Predicting Visual Fixations. Annu Rev Vis Sci 9:269– 291.

Kümmerer M, Bethge M, Wallis TSA (2022) DeepGaze III: Modeling free-viewing human scanpaths with deep learning. J Vis 22:7.

Kutas M, Federmeier KD (2011) Thirty Years and Counting: Finding Meaning in the N400 Component of the Event-Related Brain Potential (ERP). Annu Rev Psychol 62:621–647.

LeCun Y (2022) A Path Towards Autonomous Machine Intelligence Version 0.9.2, 2022-06-27. Open Rev 62:1–62.

Leszczyński M, Barczak A, Kajikawa Y, Ulbert I, Falchier AY, Tal I, Haegens S, Melloni L, Knight RT, Schroeder CE (2020) Dissociation of broadband high-frequency activity and neuronal firing in the neocortex. Sci Adv 6:eabb0977.

Leszczynski M, Bickel S, Nentwich M, Russ BE, Parra L, Lakatos P, Mehta A, Schroeder CE (2023) Saccadic modulation of neural excitability in auditory areas of the neocortex. Curr Biol 33:1185–1195.e6.

Leszczynski M, Chaieb L, Staudigl T, Enkirch SJ, Fell J, Schroeder CE (2021) Neural activity in the human anterior thalamus during natural vision. Sci Rep 11:17480.

Lindsay GW (2021) Convolutional Neural Networks as a Model of the Visual System: Past, Present, and Future. J Cogn Neurosci 33:2017–2031.

Logothetis NK, Sheinberg DL (1996) Visual object recognition. Annu Rev Neurosci 19:577–621.

Ma WJ, Beck JM, Latham PE, Pouget A (2006) Bayesian inference with probabilistic population codes. Nat Neurosci 9:1432–1438.

Madison A, Callahan-Flintoft C, Thurman SM, Hoffing RAC, Touryan J, Ries AJ (2025) Fixation-related potentials during a virtual navigation task: The influence of image statistics on early cortical processing. Atten Percept Psychophys Available at: https://link.springer.com/10.3758/s13414-024-03002-5 [Accessed February 7, 2025].

Maris E, Oostenveld R (2007) Nonparametric statistical testing of EEG- and MEG-data. J Neurosci Methods 164:177–190.

Millidge B, Seth A, Buckley CL (2022) Predictive Coding: a Theoretical and Experimental Review. Available at: http://arxiv.org/abs/2107.12979 [Accessed October 6, 2024].

Mischler G, Raghavan VS, Keshishian M, Mesgarani N (2023) naplib-python: Neural acoustic data processing and analysis tools in python. Softw Impacts 17:100541.

Mishra A, Tostaeva G, Nentwich M, Espinal E, Markowitz N, Winfield J, Freund E, Gherman S, Leszczynski M, Schroeder CE, Mehta AD, Bickel S (2025) Motifs of human high-frequency oscillations structure processing and memory of continuous audiovisual narratives. Sci Adv 11:eadv0986.

Nau M, Greene A, Tarder-Stoll H, Lossio-Ventura JA, Pereira F, Chen J, Baldassano C, Baker CI (2025) Neural and behavioral reinstatement jointly reflect retrieval of narrative events. Nat Commun 16:7865.

Nentwich M, Leszczynski M, Russ BE, Hirsch L, Markowitz N, Sapru K, Schroeder CE, Mehta AD, Bickel S, Parra LC (2023) Semantic novelty modulates neural responses to visual change across the human brain. Nat Commun 14:2910.

Nentwich M, Leszczynski M, Schroeder CE, Bickel S, Parra LC (2025) Intrinsic dynamic shapes responses to external stimulation in the human brain. eLife 14:RP104996.

Neupane S, Guitton D, Pack CC (2020) Perisaccadic remapping: What? How? Why? Rev Neurosci 31:505–520.

Nunez-Elizalde AO, Huth AG, Gallant JL (2019) Voxelwise encoding models with non-spherical multivariate normal priors. NeuroImage 197:482–492.

Otero-Millan J, Troncoso XG, Macknik SL, Serrano-Pedraza I, Martinez-Conde S (2008) Saccades and microsaccades during visual fixation, exploration, and search: Foundations for a common saccadic generator. J Vis 8:21.

Papademetris X, Jackowski MP, Rajeevan N, DiStasio M, Okuda H, Constable RT, Staib LH (2006) BioImage Suite: An integrated medical image analysis suite: An update. Insight J 2006:209.

Pazo-Alvarez P, Cadaveira F, Amenedo E (2003) MMN in the visual modality: a review. Biol Psychol 63:199–236.

Pedregosa F et al. (2011) Scikit-learn: Machine Learning in Python. Mach Learn PYTHON.

Pitcher D, Dilks DD, Saxe RR, Triantafyllou C, Kanwisher N (2011) Differential selectivity for dynamic versus static information in face-selective cortical regions. NeuroImage 56:2356–2363.

Pitcher D, Ianni G, Ungerleider LG (2019) A functional dissociation of face-, body- and scene-selective brain areas based on their response to moving and static stimuli. Sci Rep 9:8242.

Raghavan VS, O’Sullivan J, Bickel S, Mehta AD, Mesgarani N (2023) Distinct neural encoding of glimpsed and masked speech in multitalker situations Bizley JK, ed. PLOS Biol 21:e3002128.

Raghavan VS, Parra LC (2025) Neural encoding of linguistic features during natural sentence reading. iScience 28:112798.

Rajkai C, Lakatos P, Chen C-M, Pincze Z, Karmos G, Schroeder CE (2008) Transient Cortical Excitation at the Onset of Visual Fixation. Cereb Cortex 18:200–209.

Ramezani F, Kheradpisheh SR, Thorpe SJ, Ghodrati M (2019) Object categorization in visual periphery is modulated by delayed foveal noise. J Vis 19:1.

Reilly J et al. (2025) What we mean when we say semantic: Toward a multidisciplinary semantic glossary. Psychon Bull Rev 32:243–280.

Richter D, Kietzmann TC, Lange FP de (2024) High-level visual prediction errors in early visual cortex. PLOS Biol 22:e3002829.

Ries AJ, Slayback D, Touryan J (2018) The fixation-related lambda response: Effects of saccade magnitude, spatial frequency, and ocular artifact removal. Int J Psychophysiol 134:1–8.

Robinson D (2022) The function and phylogeny of eye movements. In: Progress in Brain Research, pp 1–14. Elsevier. Available at: https://linkinghub.elsevier.com/retrieve/pii/S0079612321002004 [Accessed July 30, 2025].

Roelfsema PR, Holtmaat A (2018) Control of synaptic plasticity in deep cortical networks. Nat Rev Neurosci 19:166–180.

Russ BE, Leopold DA (2015) Functional MRI mapping of dynamic visual features during natural viewing in the macaque. NeuroImage 109:84–94.

Russ BE, Petkov CI, Kwok SC, Zhu Q, Belin P, Vanduffel W, Hamed SB (2021) Common functional localizers to enhance NHP & cross-species neuroscience imaging research. NeuroImage 237:118203.

Saravanan V, Berman GJ, Sober SJ (2020) Application of the hierarchical bootstrap to multi-level data in neuroscience. Neurons Behav Data Anal Theory 3:https://nbdt.scholasticahq.com/article/13927-application-of-the-hierarchical-bootstrap-to-multi-level-data-in-neuroscience.

Schiller PH, Tehovnik EJ (2005) Neural mechanisms underlying target selection with saccadic eye movements. In: Progress in Brain Research, pp 157–171. Elsevier. Available at: https://linkinghub.elsevier.com/retrieve/pii/S0079612305490123 [Accessed May 14, 2026].

Seidlitz J, Sponheim C, Glen D, Ye FQ, Saleem KS, Leopold DA, Ungerleider L, Messinger A (2018) A population MRI brain template and analysis tools for the macaque. NeuroImage 170:121–131.

Solomon SS, Tang H, Sussman E, Kohn A (2021) Limited Evidence for Sensory Prediction Error Responses in Visual Cortex of Macaques and Humans. Cereb Cortex 31:3136–3152.

Stankov AD, Touryan J, Gordon S, Ries AJ, Ki J, Parra LC (2021) During natural viewing, neural processing of visual targets continues throughout saccades. J Vis 21:7.

Stewart EEM, Valsecchi M, Schütz AC (2020) A review of interactions between peripheral and foveal vision. J Vis 20:2.

Summerfield C, de Lange FP (2014) Expectation in perceptual decision making: neural and computational mechanisms. Nat Rev Neurosci 15:745–756.

Telesford QK, Gonzalez-Moreira E, Xu T, Tian Y, Colcombe SJ, Cloud J, Russ BE, Falchier A, Nentwich M, Madsen J, Parra LC, Schroeder CE, Milham MP, Franco AR (2023) An open-access dataset of naturalistic viewing using simultaneous EEG-fMRI. Sci Data 10:554.

Thorat S, Doerig A, Kroner A, Amme C, Kietzmann TC (2025) Predicting upcoming visual features during eye movements yields scene representations aligned with human visual cortex. Available at: http://arxiv.org/abs/2511.12715 [Accessed May 26, 2026].

Valsecchi M, Gegenfurtner KR (2016) Dynamic Re-calibration of Perceived Size in Fovea and Periphery through Predictable Size Changes. Curr Biol 26:59–63.

Vinken K, Sharma S, Livingstone MS (2025) Mapping macaque to human cortex with natural scene responses. Proc Natl Acad Sci 122:e2512619122.

Walsh KS, McGovern DP, Clark A, O’Connell RG (2020) Evaluating the neurophysiological evidence for predictive processing as a model of perception. Ann N Y Acad Sci 1464:242.

Webster MA (2015) Visual Adaptation. Annu Rev Vis Sci 1:547–567.

Weierich MR, Wright CI, Negreira A, Dickerson BC, Barrett LF (2010) Novelty as a dimension in the affective brain. NeuroImage 49:2871–2878.

Wen Z, Li Y (2021) Toward Understanding the Feature Learning Process of Self-supervised Contrastive Learning. Available at: http://arxiv.org/abs/2105.15134 [Accessed November 5, 2024].

Westerberg JA et al. (2024) Stimulus history, not expectation, drives sensory prediction errors in mammalian cortex. :2024.10.02.616378 Available at: https://www.biorxiv.org/content/10.1101/2024.10.02.616378v1 [Accessed October 7, 2024].

Wild B, Treue S (2021) Primate extrastriate cortical area MST: a gateway between sensation and cognition. J Neurophysiol 125:1851–1882.

Williams MA, Baker CI, Op de Beeck HP, Mok Shim W, Dang S, Triantafyllou C, Kanwisher N (2008) Feedback of visual object information to foveal retinotopic cortex. Nat Neurosci 11:1439–1445.

Wolf C, Schütz AC (2015) Trans-saccadic integration of peripheral and foveal feature information is close to optimal. J Vis 15:1.

Yamins DLK, DiCarlo JJ (2016) Using goal-driven deep learning models to understand sensory cortex. Nat Neurosci 19:356–365.

Yamins DLK, Hong H, Cadieu CF, Solomon EA, Seibert D, DiCarlo JJ (2014) Performance-optimized hierarchical models predict neural responses in higher visual cortex. Proc Natl Acad Sci 111:8619–8624.

Yao Y, Stebner A, Tuytelaars T, Geirnaert S, Bertrand A (2024) Identifying temporal correlations between natural single-shot videos and EEG signals. J Neural Eng 21:016018.

Yeo BTT, Krienen FM, Sepulcre J, Sabuncu MR, Lashkari D, Hollinshead M, Roffman JL, Smoller JW, Zöllei L, Polimeni JR, Fischl B, Liu H, Buckner RL (2011) The organization of the human cerebral cortex estimated by intrinsic functional connectivity. J Neurophysiol 106:1125–1165.

