## Supplementary figures and images for "During natural vision, semantic novelty modulates fixation-related processing in primate cortex"

### Figure S1

#### Clustering of Fixation Patch Feature Correlation

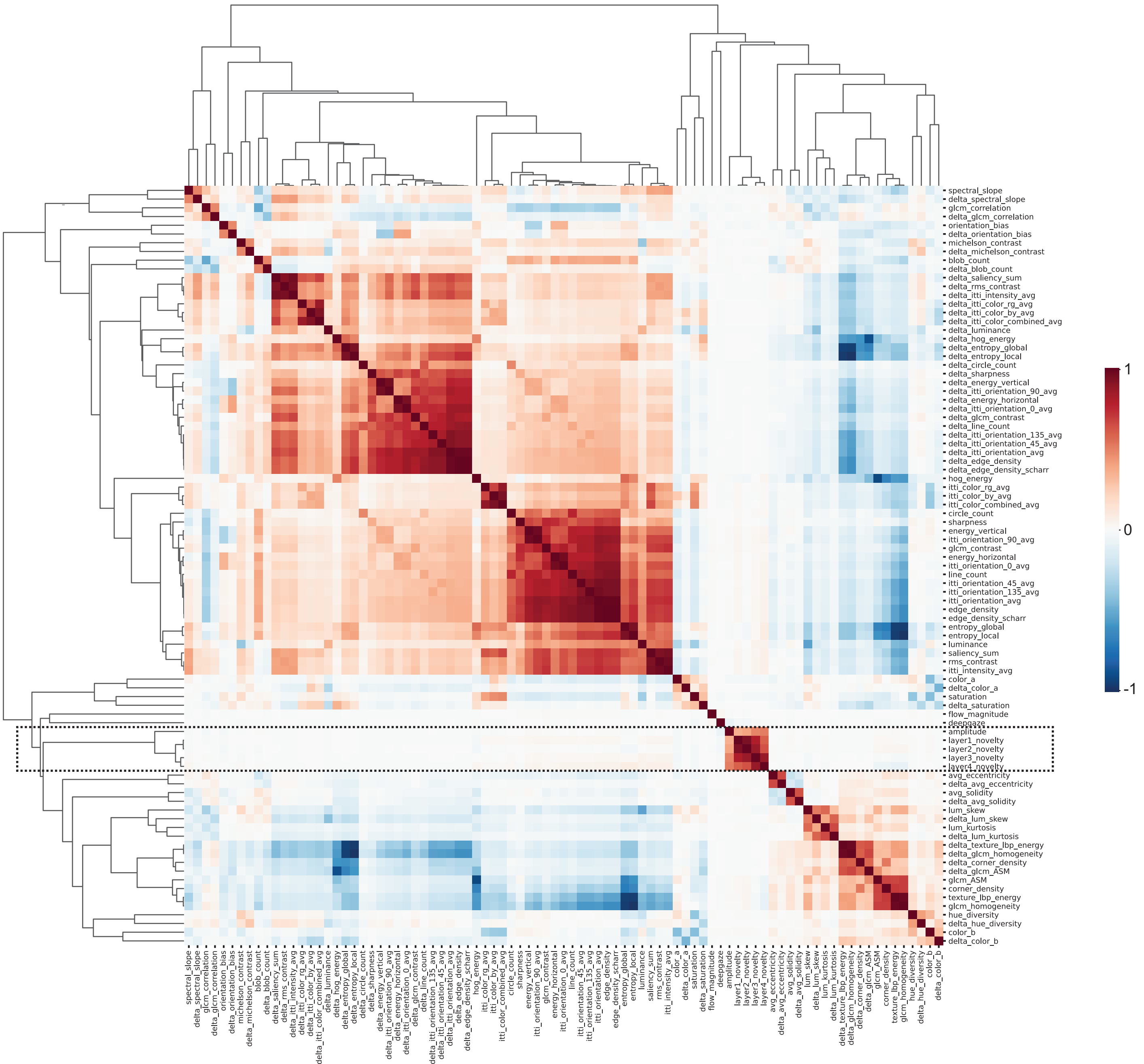

### Figure S2

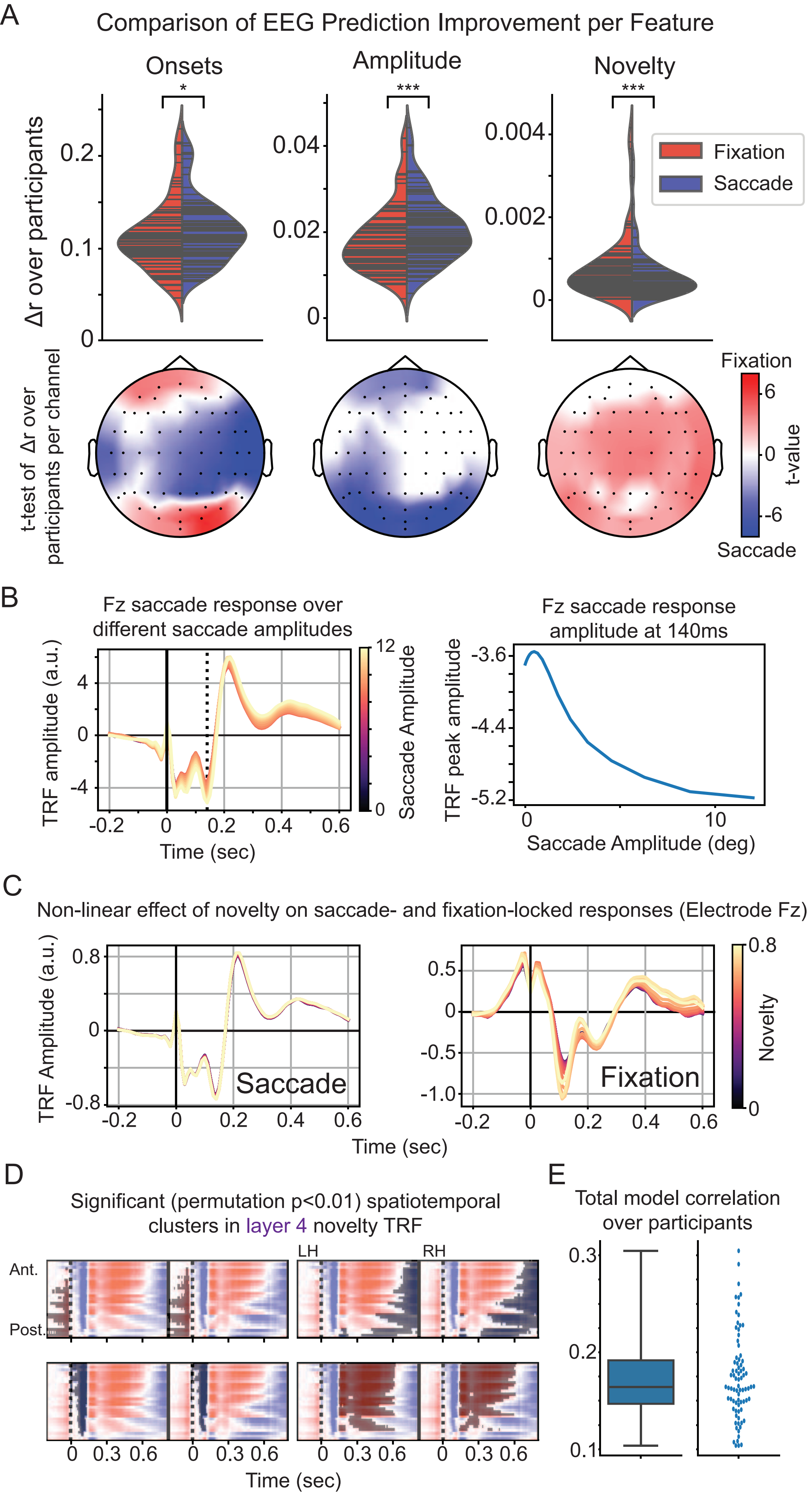

### Figure S3

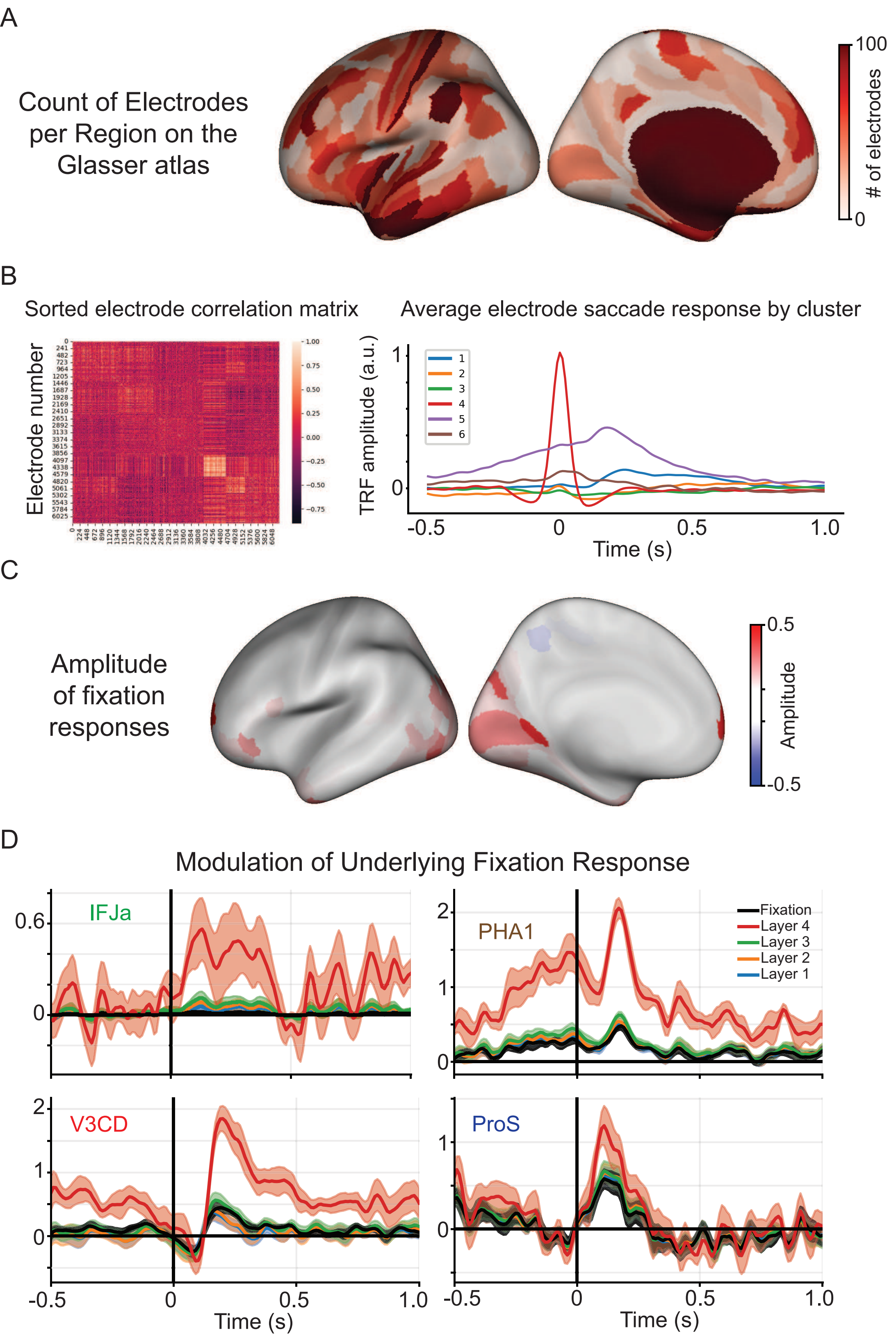
